# Effects of global change on animal biodiversity in boreal forest landscape: an assemblage dissimilarity analysis

**DOI:** 10.1101/2022.01.31.477297

**Authors:** Ilhem Bouderbala, Guillemette Labadie, Jean-Michel Béland, Yan Boulanger, Christian Hébert, Patrick Desrosiers, Antoine Allard, Daniel Fortin

## Abstract

Despite an increasing number of studies highlighting the impacts of climate change on boreal species, the main factors that will drive changes in species assemblages remain ambiguous. We quantify two climate-induced pathways based on direct and indirect effects on species occupancy and assemblage dissimilarity under different harvest management scenarios. The direct climate effects illustrate the impact of climate variables while the indirect effects are reflected through the changes in land cover composition. To understand the main causes in assemblage dissimilarity, we analyze the regional and the latitudinal species assemblage dissimilarity by decomposing it into *balanced variation in species occupancy and occurrence* and *occupancy and occurrence gradient*. We develop empirical models to predict the distribution of more than 100 bird and beetle species in the Côte-Nord region of Québec over the next century. Our results show that the two pathways are complementary and alter biodiversity, mainly caused by balanced variation in species occupancy and occurrence. At the regional scale, both effects have an impact on decreasing the number of winning species. Yet, responses are much larger in magnitude under mixed climate effects (a mixture of direct and indirect effects). Regional assemblage dissimilarity reached 0.77 and 0.69 under mixed effects versus 0.09 and 0.10 under indirect effects for beetles and birds, respectively, between RCP8.5 and baseline climate scenarios when considering harvesting. Therefore, inclusion of climatic variables considers aspects other than just those related to forest landscapes, such as life cycles of animal species. Latitudinally, assemblage dissimilarity increased following the climate conditions pattern. Our analysis contributes to the understanding of how climate change alters biodiversity by reshaping community composition and highlights the importance of climate variables in biodiversity prediction.

## 1 Introduction

Global climate warming affects the functionality of ecosystems by modifying forest composition and biomass, which in turn has repercussions for biodiversity and species assemblages (Kelly and Goulden, 2008; Hillebrand et al., 2010; Thuiller et al., 2011; Pachauri et al., 2014; Zhang and Liang, 2014). In fact, many studies have anticipated that anthropogenic radiative forcing will alter boreal biodiversity and ecosystems (Pachauri et al., 2014; Tremblay et al., 2018; Cadieux et al., 2020). For example, global scale predictions have shown that the potential high emissions of greenhouse gases would lead mainly to negative effects on biodiversity (Pachauri et al., 2014). By contrast, other results have been published anticipating an increase in biodiversity during this century, which is referred to *the northern biodiversity paradox* (Matthews et al., 2004; Morin and Thuiller, 2009; Berteaux et al., 2010). This paradox suggests that the ecological niches of some “southerly” species would increase in size due to expansion beyond the northern periphery of their ranges (Berteaux et al., 2010).

In Canada, temperature has increased by 1.7 °C since 1948, twice as fast as the global average (Bush and Lemmen, 2019). This increase in temperature could lead to the northward migration of thermophilous hardwood tree species to the detriment of boreal conifers, particularly mid-to-late-successional species (Duveneck et al., 2014; Boulanger et al., 2017; Boulanger and Puigdevall, 2021). Moreover, climate change is expected to influence wildfire activity directly (Boulanger and Puigdevall, 2021), which would favour pioneer and fire-adapted boreal tree species (Boulanger et al., 2017). Significant changes in species composition are expected within the transition zone between boreal and temperate biomes, where several tree species are currently reaching their thermal limits (Brice et al., 2020; Boulanger and Puigdevall, 2021). Global warming could also drive the occurrence of more extreme climatic events, including severe droughts (Kumar, 2013; Masson-Delmotte et al., 2018), thereby reducing the productivity of several boreal tree species through increasing metabolic respiration (Girardin et al., 2016). Climate-induced changes in insect outbreak regimes, notably those in spruce budworm (SBW) (*Choristoneura fumiferana*) can lead to severe defoliation and death of firs (*Abies* spp.) and spruces (*Picea* spp.), and alter forest successional pathways (Pureswaran et al., 2015; Labadie et al., 2021).

Global change might influence biodiversity through behavioral, morphological and physiological changes of organisms or might even influence gene flow (Naeem et al., 2012; Wisz et al., 2013; Sergio et al., 2018; Matuoka et al., 2020; Boulanger and Puigdevall, 2021; Micheletti et al., 2021). The implications of global change on biodiversity could be synthesized through direct and indirect effects. On one hand, the indirect effects are associated with habitat change (e.g., vegetation) that could occur through various events such as natural and anthropogenic disturbances (Boulanger et al., 2017). On the other hand, direct effects are mainly characterized by the influence of climate and meteorological conditions on the organisms (Micheletti et al., 2021). Both types of effects should be considered when studying the impacts of global warming on ecosystem functioning. For example, ecosystem services can depend on the magnitude of both direct and indirect pathways on global biodiversity. Understanding both types of effects should thus help identify the best conservation actions that are required to maintain valued services. If habitat-based global change is projected to decrease species occupancy, land management actions could be adapted by targeting vegetation restoration. Otherwise, species translocation actions could be adopted if direct changes are anticipated that would cause a decline in species occupancy (Micheletti et al., 2021).

Climate and land-use changes may interact in complex ways to impact changes in wildlife habitats (Bentz et al., 2010; Tremblay et al., 2018). For example, an increase in disturbance rate due to warmer conditions could favor the regeneration of warm-adapted, pioneer tree species (Brice et al., 2020; Boulanger and Puigdevall, 2021), which could affect biodiversity through adjustments in species ecological traits. Moreover, the synergistic effects of harvest and natural disturbances with direct climate change could alter species assemblages, and, therefore, biodiversity. In this work, we investigated future climate-induced variations in bird and beetle assemblages in Québec’s boreal forest under different forest harvesting scenarios. This will enhance our knowledge regarding the interactive effects of climate change and forest use on future biodiversity. We assessed effects of future climate conditions on these assemblages by comparing forest landscapes that were simulated under two anthropogenic radiative-forcing scenarios, i.e., Representative Concentration Pathway (RCP) 4.5 and RCP 8.5 (van Vuuren et al., 2011) with landscapes that are simulated under average historical (baseline) climate conditions. Forest landscapes were simulated through a spatially explicit raster-based forest landscape model to capture key forest ecosystem processes. We further assessed how forest management may affect future assemblages by simulating species occupancy if current harvest levels are maintained vs there would be no harvest activities. We analyze the effects of global change in two ways: (1) indirectly, i.e., habitat-based change (Wisz et al., 2013; Boulanger and Puigdevall, 2021; Micheletti et al., 2021) and (2) through a mixed effect combining indirect and direct effects, i.e., those stemming from climate variables *per se* (Thuiller et al., 2018). The choice of the two pathways was made to quantify separately their implications on biodiversity in a cascading configuration. More precisely, starting first with habitat effects, through Habitat-Only-Based Models (HOBMs), and adding afterward meteorological conditions, through Climate-Habitat-Based Models (CHBMs). We also determined the main drivers of assemblage dissimilarity following the two pathways.

We opted for a species distribution modeling (SDM) approach (Guisan and Zimmermann, 2000; Guisan and Thuiller, 2005; Peterson et al., 2011) to model the single-species occurrence probability based on extensive field surveys of beetles and birds. We analyzed the assemblage structure based on continuous occurrence probabilities (probability-based) (Guisan and Thuiller, 2005; Grenié et al., 2020) to avoid overprediction risks (Gelfand et al., 2005; Dubuis et al., 2011; Grenié et al., 2020). It has been previously demonstrated that probability-based richness provided a better fit of the actual richness than would the threshold-based approach (Grenié et al., 2020). We computed dissimilarity measures (Baselga, 2010; Albouy et al., 2012; Baselga, 2013) between different climate scenarios under the two harvesting levels. In this context, we adapted continuous-based decomposition in the context of occurrence probabilities to detail the two components of *Beta*-*diversity* : *balanced variation in species abundances* and *abundance gradient*, which are generalizations of turnover and nestedness (Baselga, 2013). Quantifying the main causes of change in biodiversity could be very helpful in assessing the potential underlying determinants because species replacement and nestedness, for example, are two different ways of generating assemblage dissimilarity (Baselga, 2013).

## 2 Material and methods

### 2.1 Study area and occurrence data

The study area is located in the Côte-Nord region of Québec, Canada (48°*N* to 53°*N*, 65°*W* to 71°*W*), within an area of 114118 km^2^ (Fig. 1). The northern part of the study area belongs to the black spruce-feather moss bioclimatic domain, and is dominated by black spruce (*Picea mariana* [Mill.] BSP) and balsam fir (*Abies balsamea* [L.] Miller). Wildfires are the major natural disturbances in northern areas that have yet to be logged (Boucher et al., 2017). The southern part of the study area belongs to the eastern balsam fir - white birch (*Betula papyrifera* Marsha.) subdomain, mostly dominated by balsam fir and white spruce (*Picea glauca* [Moench] Voss) mixed with white (paper) birch. Forest harvesting had been the main source of forest disturbance since the late 1990s in this latter area (Bouchard and Pothier, 2011). In Québec, logging affected around 0.8% of public forest annually (Bureau du Forestier en chef, 2010). This part of the territory also has been affected by an outbreak of the SBW that began in 2006 and which is still ongoing. Tree mortality began around 2015 and has subsequently increased.

**Figure 1:**
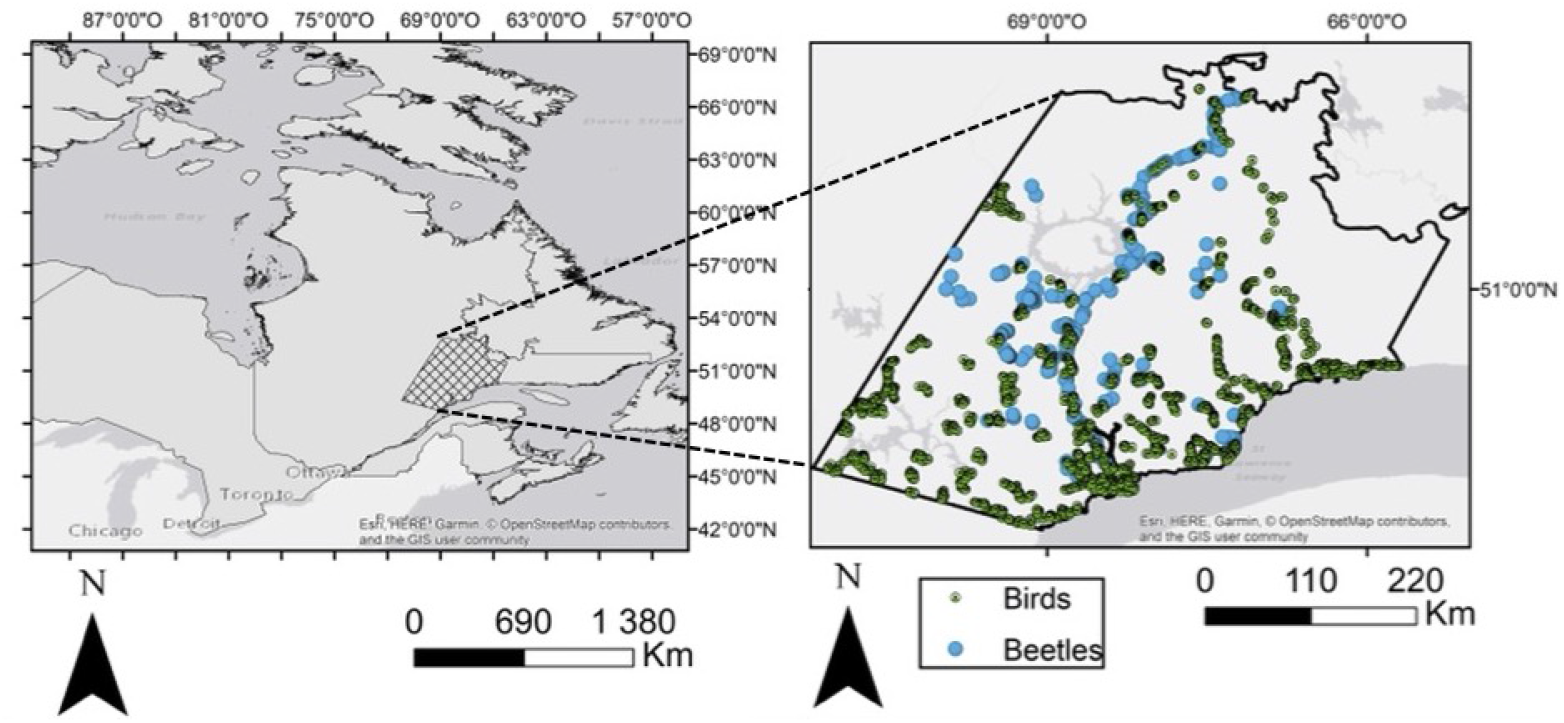
Study area with the presence-absence distribution of the two taxa.

We used presence-absence data that were collected between 2004 and 2018 to model species distributions. Given that we were mainly focusing on the impacts of fire and harvesting, we wanted to remove sites that were located in stands heavily damaged by the SBW outbreak (Ministère des Forêts de la Faune et des Parcs, 2018) by using a cumulative index of defoliation severity from 2007 to 2018 (Labadie et al., 2021). Annual defoliation severity was based on aerial surveys characterizing damage that was incurred by SBW since 2006 (Ministère des Forêts de la Faune et des Parcs, 2018) and was classified between 0 and 3, with 3 indicating the highest level of defoliation. The cumulative severity of the outbreak was obtained by summing the annual severity in Labadie et al. (2021). Sites with cumulative severity values of 10 and above were discarded from analyses. This threshold attests for a strong impact of the SBW over a long period, which modifies stand structure and therefore affects species presence. Indeed, Labadie et al. (2021) showed (see Fig. 2 in this paper) that, at this level, the response of shrubs is clearly stronger that of herbaceous plants (no overlap in the 95% CI), which indicates important changes in forest structure that modify habitat of many species.

**Figure 2:**
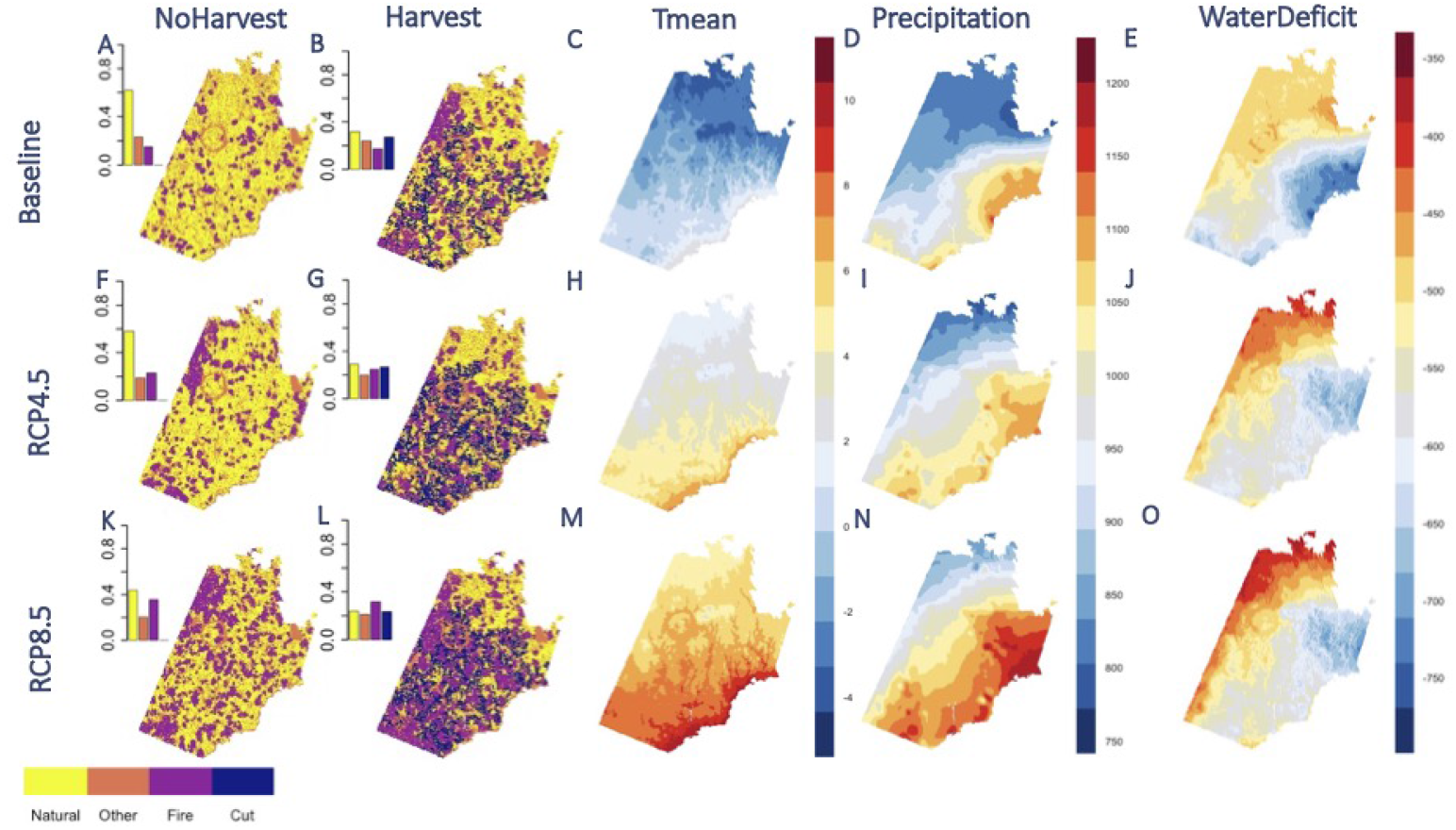
The distribution of the land cover and three climate variables under the simulated management scenarios in 2100. **A-B, F-G** and **K-L** represent the distribution of land cover over the six study scenarios that were classified into four large cover classes: *Natural*, which included conifer dense, conifer open, mixed wood and open habitat; *Fire* and *Cut* for the land cover disturbed by fire and Harvest; and *Others* for the rest. The barplots represent the frequency of each class in the map. **D-F, J-L** and **P-R** represent the distribution of mean temperature, precipitation, and water deficit for the three climate scenarios **Baseline, RCP4.5** and **RCP8.5**, respectively.

We used the data from the Atlas of Breeding Birds (Atlas des oiseaux nicheurs du Québec, 2018), which were based on species occurrences that were detected using unlimited distance 5-minute point counts (Bibby et al., 2000), which were collected during the breeding season (late May to mid-July) between 2010 and 2018. For beetles, we merged different databases that had been collected in 2004, 2005, 2007 and 2011 (Janssen et al., 2009; Légaré et al., 2011; Bichet et al., 2016). In addition, we used data from 54 sites that had been sampled in 2018 in the northern part of the territory, along the northeast principal road going to Labrador. The sampling protocols were characterized by one multidirectional flight-interception trap per site, which sampled flying beetles, and four meshed pitfall traps per site, which sampled epigeic beetles during their peak period of activity (June-August) (Janssen et al., 2009; Bichet et al., 2016). For beetles, we used species-level identifications wherever possible; otherwise, we standardized the identification to the genus level (around 92% initial identifications were considered at the species level).

### 2.2 Predictor variables

To predict species occurrence, we used climate and land cover variables that were grouped in two classes of models: Climate-Habitat-Based (CHBMs) and Habitat-Only-Based (HOBMs). These two model classes were designed to study climate-induced effects, as follows: CHBMs for the mixed effects; and HOBMs for the indirect climate effects. Initially, we generated 22 potential climate variables at a 250-m resolution using the BioSim platform (Régnière et al., 2017), including the annual average of temperature, precipitation and water deficit between 2004 and 2018 (see Table. 1 in Supporting Information for the description of all potential predictor variables). BioSIM simulates daily maximum and minimum temperatures, precipitation, water deficit, mean daily relative humidity and wind speed by matching georeferenced sources of daily weather data to spatially georeferenced points. BioSIM uses spatial regression to adjust weather data for differences in latitude, longitude, and elevation between the sources of weather data and each field location (for more details, see (Boulanger et al., 2018a)). In our case, the spatially referenced points were 15 000 points that were randomly located across the entire Province of Québec, whereas weather data were daily data originating from discrete weather stations that were located within the province. We generated climate variables at a 250 m scale by spatially interpolating data from the 15 000 random points using kriging and elevation as a drifting variable. Land cover maps from the Canadian National Forest Inventory (NFI) were used to generate land cover variables based on k-nearest-neighbor interpolation at a 250-m resolution that was referenced to the year 2001 (Beaudoin et al., 2014). To estimate forest composition, we used the relative importance of tree species groups (conifer and deciduous species), treed land and tree canopy-closure maps from these NFI data to generate five natural land cover classes: closed-canopy conifer forest; open-canopy mature conifer forest; mixed forest; open area; and others (Labadie et al., 2021) (for more details, see Table 2 in Supporting Information). We used four of natural land cover categories together with six disturbance classes. For the disturbed stands (by fire or harvest), we subdivided them into three age-classes: [0, 10], [10, 20] and [20, 50] years. In addition, we considered stand age and the distance to the nearest burned area as potential predictor variables. Stand age maps were based on the year 2001 (Beaudoin et al., 2014) with an update according to the registered fire and harvest disturbances between 2004 and 2018. Furthermore, it has been mentioned in Bichet et al. (2016) and Zhao et al. (2013) that the influence of the landscape varied between 400 m for beetles to 1000 m for birds. Consequently, we used a matrix of 20 pixels centred on the focal pixel (21 pixels in total) to calculate frequencies of the land cover (see Table.1 and Table. 2 in Supporting Information for descriptions of the 10 land cover variables that were used in the study).

### 2.3 Climate and forest management scenarios

We obtained future climate projections from the Canadian Earth System Model version 2 (CanESM2) for RCP 4.5 and RCP 8.5 for the period 2071-2100, which were further downscaled to a 10-km resolution using ANUSPLIN (McKenney et al., 2013). Future monthly normals at each random point that was previously used to assess baseline climate were directly assessed from changes that were observed between the 1981-2010 period and future projections in the 10-km cell in which the random point was located. Daily time series were stochastically generated for each random point from these future monthly normals using BioSIM. Future climate variables at each random point were calculated by averaging these daily values from 30 BioSIM simulations (Boulanger et al., 2018a). Climate scenarios varied depending upon the mean annual temperature that was expected to increase between 3 °C (RCP4.5) and 7.5 °C (RCP8.5) throughout the southern boreal region by 2100 (compared to 2000, see Fig. 2), while average precipitation was projected to increase between 7% and 10% regionally, with relatively small differences among scenarios (Boulanger et al. (2018b); Boulanger and Puigdevall (2021), see Fig. 2).

The forest landscapes were simulated using the spatially explicit raster-based forest landscape model LANDIS-II (Scheller and Mladenoff, 2004), which simulates stand- and landscape-scale processes, including forest succession, seed dispersal and natural (wildfires and spruce budworm outbreaks) and anthropogenic (harvest) disturbances. This model has been extensively used in Québec over the last decade and has been thoroughly validated under various forest conditions (Boulanger et al., 2017; Taylor et al., 2017; Tremblay et al., 2018; Boulanger and Puigdevall, 2021). Forest landscapes were initialized for the year 2000 conditions using the NFI attribute maps (Beaudoin et al., 2014) and provincial sample plots. Tree growth and regeneration as well as wildfires were climate-sensitive in simulations. We simulated two levels of harvesting scenarios according to a gradient of harvesting pressure, from no harvesting to high intensity harvesting, similar to current management practices in Québec (harvest applied to 8% of the harvestable upland area per 10 years). The Biomass Harvest extension was used to simulate forest harvests (Gustafson et al., 2000). Only stands that contained tree cohorts that were greater than 60-years-old were allowed to be harvested. Mean harvested patch size varied between 40 km^2^ and 150 km^2^. Harvest rates were held constant throughout the simulations unless sufficient numbers of stands did not qualify for harvest. In the latter case, harvesting continued proceed until stands were no longer available. Simulations were performed from 2000 to 2100 using a 10-yr time step. Climate sensitive parameters for simulations that were performed under RCP 4.5 and RCP 8.5 were set to change in 2011-2020, 2041-2050 and 2071-2080. Forest succession, wildfire, SBW outbreaks and harvesting were simulated using Biomass Succession v5.0, Base Fire v4.0, BDA v4.0 and the Biomass Harvest v5.0 extensions, respectively. Many more details regarding LANDIS-II simulations that were performed in this study can be found in Labadie et al. (2022) as well as in Boulanger et al. (2017) and Tremblay et al. (2018).

### 2.4 The modelling strategy

We used two classes of models, i.e., Habitat-Only-Based models (HOBMs) and Climate-Habitat-Based models (CHBMs), to study indirect climate effects and mixed climate effects, respectively. The mixed effects represented a combination of direct and indirect effects (see filled arrows in Fig. 3). We divided our modeling strategy into five steps (see Fig. 4). We aimed to study the effect of climate change under different forest harvesting scenarios. Therefore, we compared baseline climate scenario with RCP 4.5 and RCP 8.5 under two forest harvesting scenarios (NoHarvest and Harvest), hence the use of two baseline scenarios (see Step 5 in Fig. 4).

**Figure 3:**
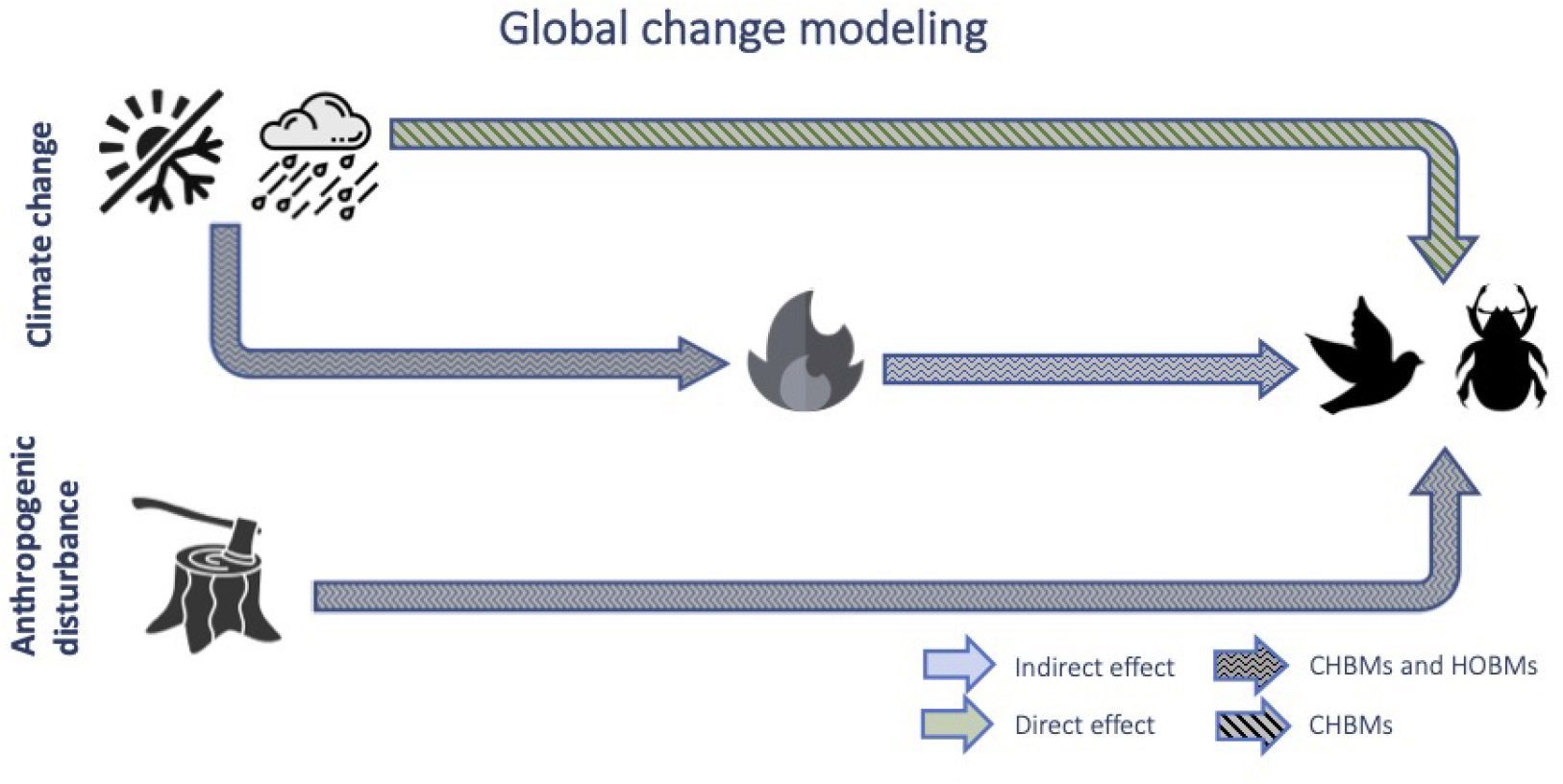
Framework used for the study. The indirect effects were generated through the change in forest composition and the direct effects included the effects of temperature, precipitation and water deficit. Abbreviations: Climate-Habitat-Based Models (CHBMs), Habitat-Only-Based Models (HOBMs).

**Figure 4:**
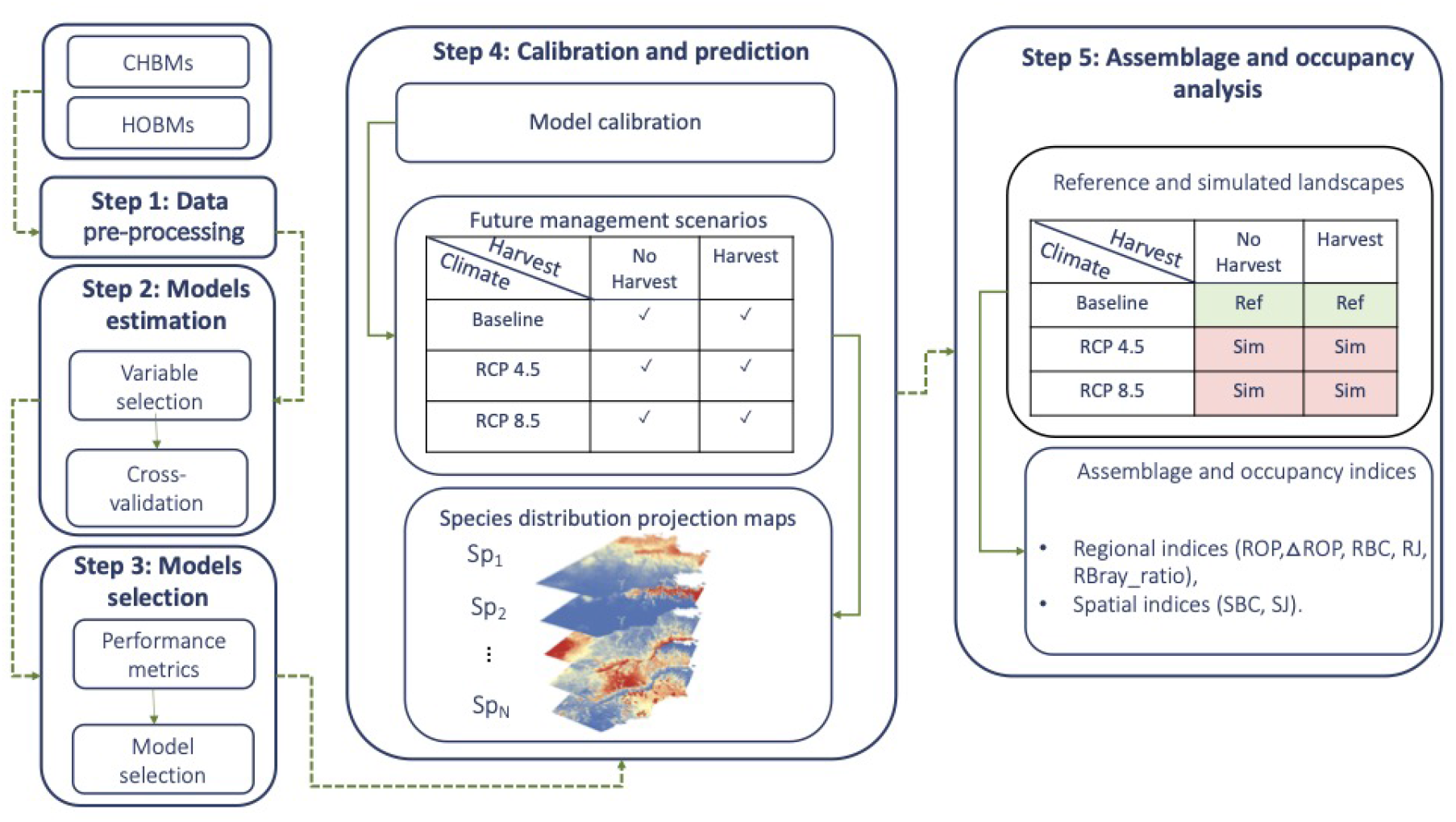
The modeling strategy that was used under Habitat-Only-Based models (HOBMs) and ClimateHabitat-Based models (CHBMs). Step 1 was dedicated to database preparation before starting the cross-validation. In Step 2, we performed the cross-validation and computed the performance metrics in Step 3. In Step 4, we calibrated SDMs only for the selected species *AUC* ≥0.7 and computed the prediction maps for the six specified scenarios. The last step was devoted to the assemblage analysis.

The modelling procedure that is described here was repeated for the two model classes (CHBMs and HOBMs) according to potential predictor variables under each class. We standardized the variables to facilitate model convergence (MacKenzie et al., 2017) prior to model calibration that was based on the corresponding database. Steps 2 and 3 correspond to a cross-validation loop, in which we split the date into 10 folds of relatively equal size, so that at each step, 9 folds were used for training and the one remaining fold was used for testing.

#### 2.4.1 Step 1: Data pre-processing

We used the following procedure for preparing the two databases: (1) we removed all of the sites that were strongly affected by the spruce budworm outbreak; and (2) we included only the more common species, with a minimum record of 1% and 5% presence among sites for birds and beetles, respectively. These percentages made sure that we only modeled species that occurred at *≥* 24 sites for birds and *≥* 14 sites for beetles.

#### 2.4.2 Step 2: Model estimation

First, we removed the highly correlated variables, based on pairwise Pearson correlation coefficient (*r*), and retained the 5 most important predictor variables (Zurell et al., 2020). To do so, we fitted a univariate Generalized linear model (GLM) with linear and quadratic terms for each variable; we ranked the predictors according to their importance, using the Akaike information criterion (AIC); we removed the highly correlated variables (|*r*| *>* 0.60). To reduce the separation in the regressions, we removed any predictor with a standard deviation value greater than 50 through a stepwise procedure.

We started with a generalized linear mixed model (GLMM) (package ‘lme4’, (Bates et al., 2015)) with a random intercept to account for differences between sampling years. We developed six and three full potential models differing only in the fixed effects for CHBMs and HOBMs model classes, respectively (see Table 3 in Supporting Information). We used interaction terms between the best temperature variable (where the AIC criterion of the corresponding univariate regression is the lowest among all selected temperature variables) with distance to the nearest burned stand and with stand age variables to include the effect of latitudinal variation.

The best models were selected based on each full-model under 10-fold cross-validation by minimizing the AIC criterion (package ‘MuMIn’, (Barton, 2015)) under the following conditions: (1) limiting the number of terms in the model between 1 and ⌈*min*(*N*_*Pre*_, *N*_*Abs*_)*/*5⌉ without counting the intercept by specifying 1 in 5 rule, where *N*_*Pre*_ and *N*_*Abs*_ represented the number of respective presence and absence records, to avoid overparameterized models; and (2) respecting the principle of marginality when the interaction terms were included in the model. For each species and for each test dataset, we calculated the occurrence probabilities matrix for the calibrated models. In total, we retained six models for CHBMs and three for the HOBMs.

#### 2.4.3 Step 3: Model selection

Once the occurrence probabilities were computed for the complete dataset for each species, we calculated the following performance metrics: (1) specificity; (2) sensitivity; (3) the area under the curve *AUC*; and (4) the true skill statistic *TSS* (Araújo et al., 2005; Allouche et al., 2006). The package ‘AUC’ (Ballings and Van den Poel, 2013) was used to calculate the Receiver Operating Characteristic (ROC) and the AUC; we used the package ‘PresenceAbsence’ (Freeman and Freeman, 2012) for the other performance metrics. Species were excluded if no model yielded *AUC* ≥ 0.7 (Araújo et al., 2005; Hosmer et al., 2013). For the selected species, we used the model with the highest AUC for projections.

#### 2.4.4 Step 4: Calibration and prediction

We estimated the parameters of the final model for the selected species based on the full dataset using the same procedure that was described for cross-validation. We subsequently used the simulated predictor variable maps for the six study scenarios and computed the occurrence probability maps.

#### 2.4.5 Step 5: Assemblage and occupancy analysis

We used the following indices to compare species assemblages between scenarios:

##### Regional occupancy probability (*ROP*)

ROP was calculated as the regional average of the occurrence probabilities for the study area (Bichet et al., 2016). We also used the percentage of change in ROP between the reference (*Ref*) and the simulated (*Sim*) scenarios (see step 5 in Fig. 4 for the definitions of *Ref* and *Sim* scenarios): 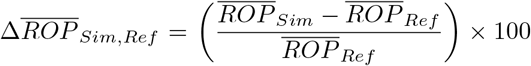, where 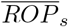 represented the average of the ROP over all species under the scenario *s* given by:

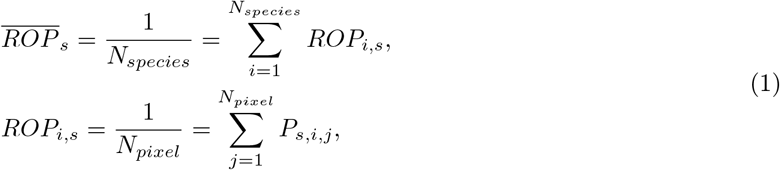

where *N*_*species*_, *N*_*pixel*_ and *P*_*s,i,j*_ represented respectively the species number, the number of pixels in the study area and the occurrence probability of the species *i* for the scenario *s* at the cell *j*.

##### Dissimilarity measures

Jaccard Index of Dissimilarity was used to assess climate change effects on species assemblage (Baselga, 2010, 2013; Legendre, 2014; Bichet et al., 2016; Barton et al., 2016; Belliard et al., 2018; Scherrer et al., 2020). We computed two different Jaccard indices, both based on Bray-Curtis dissimilarity measure, regional (*RJ*) and spatial (*SJ*) Jaccard indices computed as follows:

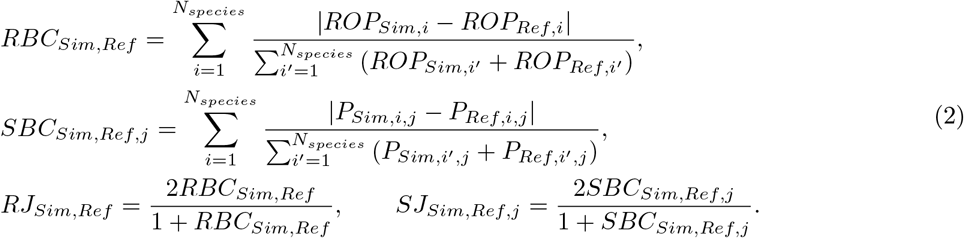

*RBC*_*Sim,Ref*_ and *SBC*_*Sim,Ref,j*_ represented respectively the regional Bray-Curtis dissimilarity between the scenarios *Sim* and *Ref*, the spatial Bray-Curtis at the cell *j* selected randomly.

From Fig. 5, we compared the performance of the Jaccard index that was based on our continuous output (*SJ*_*OP*_) and the traditional Jaccard index that was based on the presence-absence transformation (*SJ*_*Incidence*_). From the simulation results, the *SJ*_*OP*_ yields results that were close to *SJ*_*Incidence*_; moreover, performance increased with the number of species that were analyzed (see Fig. 5I). Furthermore, we added two situations with weaker binarization (Bad and Medium cases in Fig. 5) to illustrate that a gap can be generated between the two indices that is mainly due mainly to binarization error.

**Figure 5:**
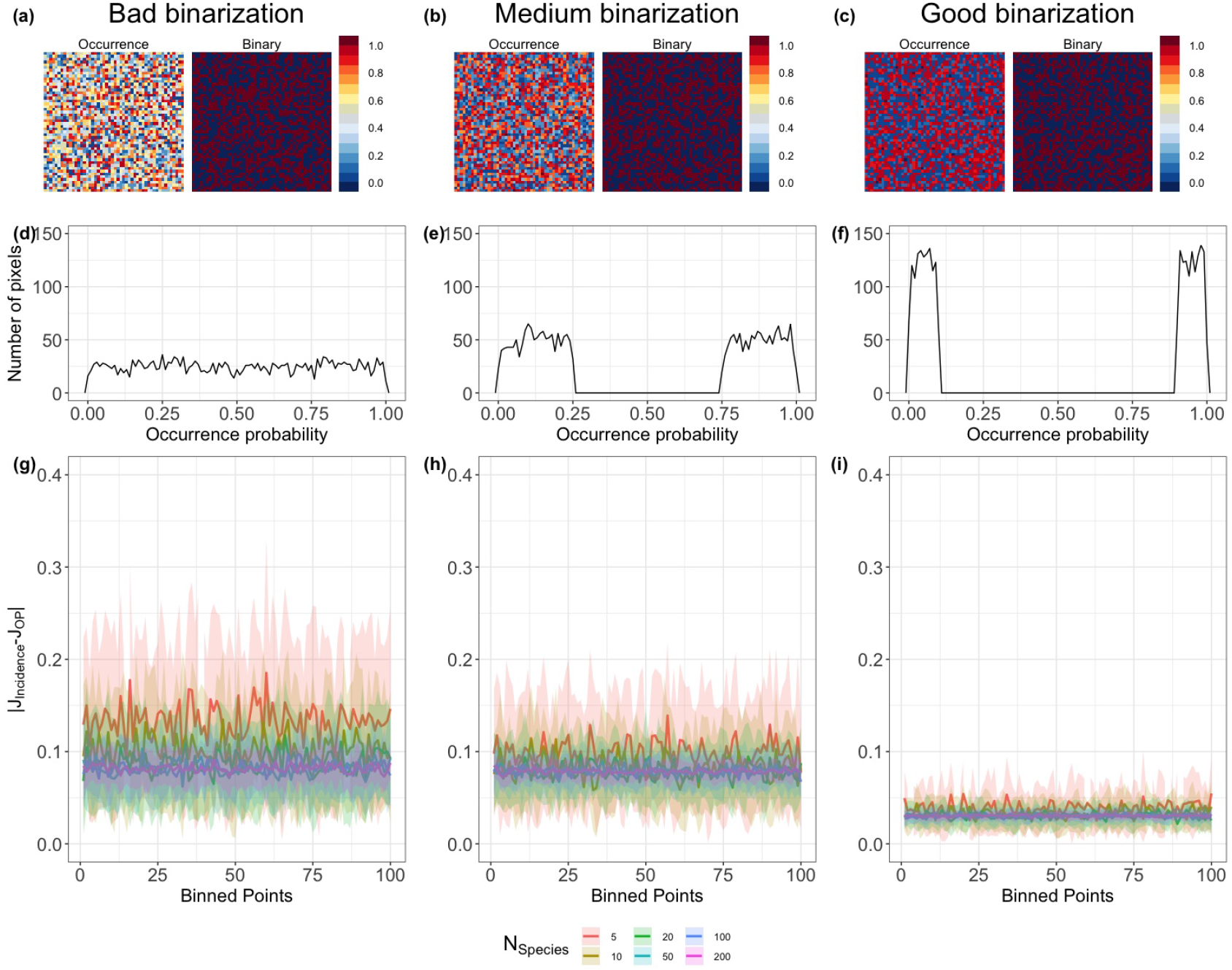
A generic Incidence-based vs. Occurrence Probabilities Jaccard dissimilarity. The three levels of binarization concerned the rank choice of the occurrence probabilities that were simulated from a mixture of uniform distributions under constraints. **(a)-(b)-(c)** represented the three simulated occurrence probabilities and the corresponding binarized maps with a 0.5 threshold for one species and under one scenario. **(d)-(e)-(f)** showed the frequency of the occurrence probability at the different pixels. **(g)-(h)-(i)** concerned the absolute difference between Jaccard dissimilarity based on the binarized data and the continuous version by varying the number of species.

##### Beta ratio

We separated the Bray-Curtis dissimilarity index into two additive components main drivers in assemblage dissimilarity: (1) the occurrence gradient (*BC*_*grad*); and (2) balanced variation in species occurrence (*BC*_*bal*). We used these notations instead of the abundance gradient and balanced variation in species abundance (Baselga, 2013), given that we worked with occurrence and occupancy probabilities. We presented a detailed explanation for each component, as follows:

- If *BC*_*grad*_*Sim,Ref*_ = 0 (*BC*_*Sim,Ref*_ = *BC*_*bal*_*Sim,Ref*_), means a **total** absence of differences in occurrences between the two scenarios. Furthermore, if *BC*_*Sim,Ref*_ ≃ 0, the assemblage structure remained almost the same. If *BC*_*Sim,Ref*_ ≃ 1, the occurrence of species in one scenario was almost perfectly balanced by the occurrence of species in the other scenario.
- If *BC*_*bal*_*Sim,Ref*_ = 0 (*BC*_*Sim,Ref*_ = *BC*_*grad*_*Sim,Ref*_), this means that all species occurrence changes from one scenario to the next were in **the same direction**. If *BC*_*Sim,Ref*_ ≃ 1, this implies that the difference in occurrence between the two scenarios is too large and in the same direction in almost all species.

To measure the fraction of each component of Bray-Curtis dissimilarity, we used the *BC*_*ratio* given by 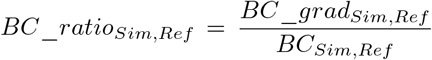. If *BC*_*ratio* > 0.5, the assisted community change was caused mostly by the occurrence gradient, whereas a value smaller than 0.5 indicated a dominance of balanced change in occurrence in the assemblage dissimilarity (Albouy et al., 2012). We used the package ‘betapart’ (Baselga and Orme, 2012) for the assemblage analysis.

The *BC*_*ratio* and its components were illustrated by using occurrence probabilities. The same formalism was adapted regionally through the regional occupancy probabilities.

## 3 Results

Of 231 candidate species of birds and beetles, 127 and 108 species were selected for projection (with *AUC* ≥ 0.7) respectively for CHBMs and HOBMs, with an average of AUC between 0.76 (±0.07) and 0.79 (±0.08), and TSS between 0.43 (±0.13) and 0.53 (±0.16) (mean ± SD). For bird species, *stand age* was the most frequently selected predictor variable with 43.9% and 71% of selection in CHBMs and HOBMs, respectively (see Fig. 6). For beetle species, mean temperature of the warmest month (*WarmMTmean*) and *stand age* were the most frequently selected variables under CHBMs and HOBMs with 33.7% and 40.3% of selection, respectively (see Fig. 6).

**Figure 6:**
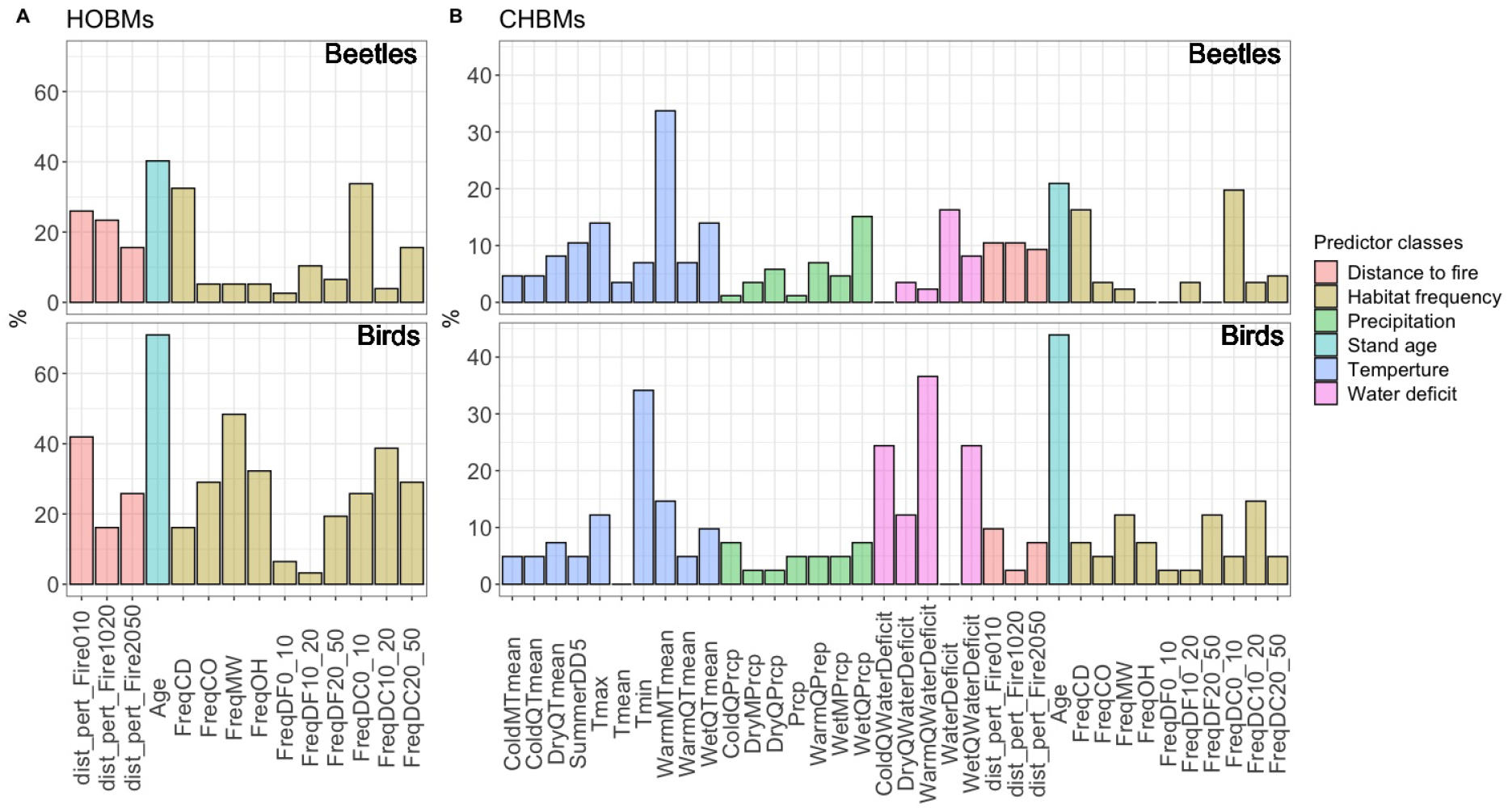
Percentage of the predictor variables that were included in the species regressions for the two taxa (See Table. 1 in Supporting Information for the variable descriptions), **A**. for Habitat-Only-Based-Models (HOBMs) and **B**. for Climate-Habitat-Based Models (CHBMs). Land cover abbreviations: Conifer Dense (CD), Conifer Open (CO), Mixed Wood (MW), Open Habitat (OH), Disturbance by Fire (DF), and Disturbance by Cut (DC). For example, under CHBMs, 21% and 44% of beetle and bird species, respectively, selected Stand age variable.

### 3.1 Occurrence and regional occupancy

The analysis of the occurrence and occupancy helped us to evaluate the direction and magnitude of potential community changes following global change. The difference between the indirect and mixed climate effects was visualized through 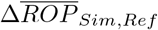 (Fig. 7A). The magnitude of changes in occupancy compared to the baseline reference scenario was larger when we included the climate variables (CHBMs) when compared to the models including only habitat variables (HOBMs). The change in occupancy was slightly more significant for birds, compared to beetles under HOBMs. Under CHBMs, we observed that occupancy increased for birds but decreased for beetles when comparing the same radiative-forcing scenarios (RCP4.5 and RCP8.5) to the baseline (the projection maps also demonstrated this result; compare Fig. 8E to Fig. 8F and Fig. 8A to Fig. 8B). However, we observed a decrease in the percentage of winner species with climate change for the two taxa, except for birds under HOBMs with no harvest (Fig. 11; see Fig. 1 in Supporting Information responses of five variable classes on eight winner and loser species). Moreover, to observe the change at the species level, Table 5 to Table 9 in the Supporting Information contained the percentage of change in the regional occupancy probability for the accepted species under the two model classes (CHBMs and HOBMs).

**Figure 7:**
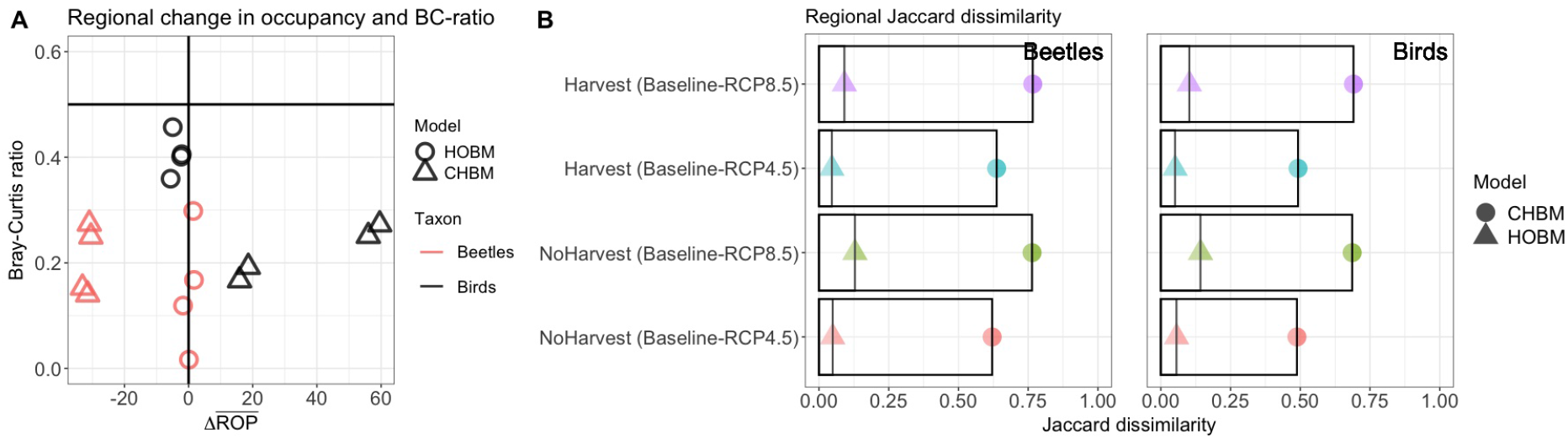
Regional dissimilarity and change in occupancy. **<u>A</u>**<u>. B</u>ray-Curtis ratio (*BC*_*ratio*) with the percentage of change in the regional occupancy probability 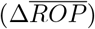 under the four compared landscapes. **B** Bird and Beetle Jaccard dissimilarity measures. Abbreviations: Climate-Habitat-Based Model (**CHBM**), Habitat-Only-Based Model (**HOBM**).

**Figure 8:**
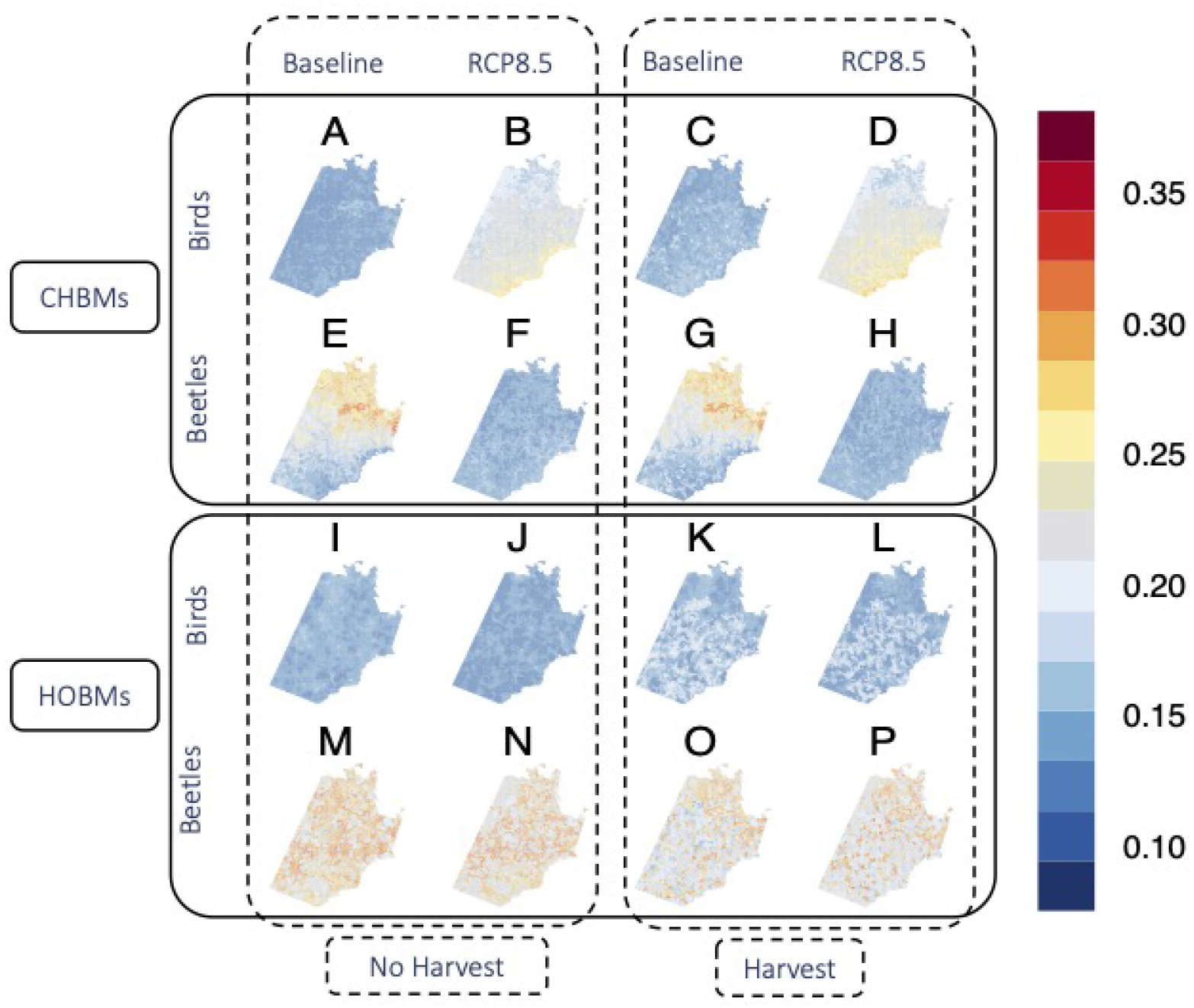
The average potential occurrence maps for each taxon according to two model classes (CHBMs and HOBMs) and four scenarios: BaselineNoHarvest (**A, E, I, M**), RCP8.5NoHarvest (**B, F, J, N**), BaselineHarvest (**C, G, K, O**) and RCP8.5Harvest (**D, H, J, P**).

### 3.2 Regional species assemblage change

An increase in assemblage dissimilarity was observed when comparing RCP4.5 and RCP8.5 to the baseline for both taxa. Based on CHBMs, regional dissimilarity from RCP4.5 to RCP8.5 increased respectively by 0.20 and 0.14 for birds and beetles under no harvest (Fig. 7B-C). Also, based on HOBMs, regional dissimilarity from RCP4.5 to RCP8.5 increased respectively by 0.07 and 0.08 for birds and beetles under no harvest. This regional dissimilarity was mainly incurred through balanced variation in species occupancy for both taxa (*BC*_*ratio* < 0.5) (see Fig. 7A).

We also observed an increase in assemblage dissimilarity from HOBMs to CHBMs for both taxa. Inclusion of the climate variables reshaped assemblage structure strongly under the two forest management levels (see differences in the Jaccard dissimilarity index between HOBMs and CHBMs, Fig. 7B-C).

### 3.3 Latitudinal changes in species assemblage

The inclusion of climate variables produced a latitudinal gradient in projections of assemblage dissimilarity. We used CHBMs to analyze how latitudinal changes in temperature and other climate variables would affect assemblage structure. A clear increasing pattern was observed in assemblage dissimilarity heading north, especially for beetle species. For birds, a slight increase in dissimilarity was observed, compared to that of beetles (see the difference between Fig. 9A and Fig. 9C).

**Figure 9:**
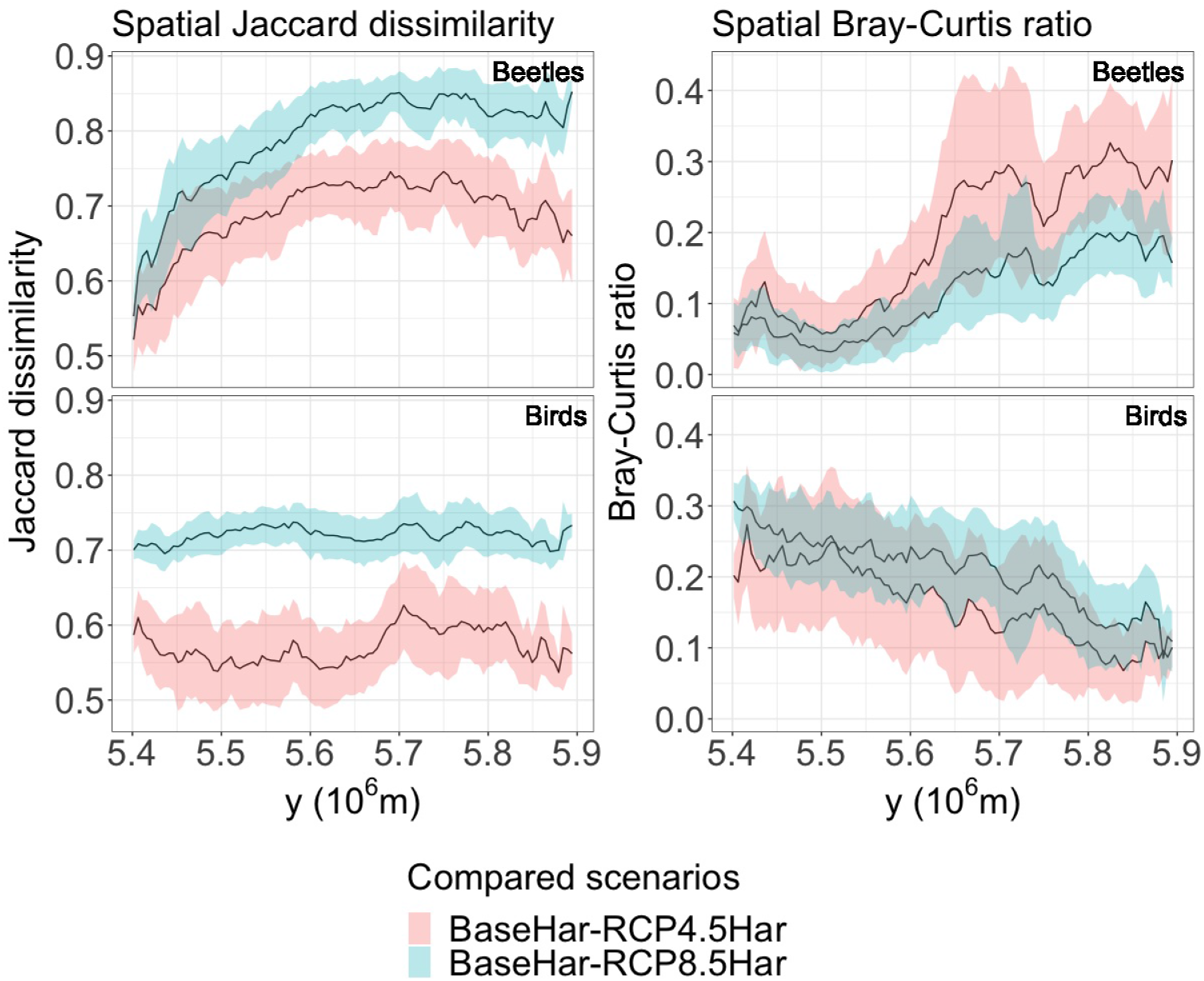
Spatial analysis under CHBMs. **A, C**. Latitudinal variation of spatial Jaccard dissimilarity of **RCP4.5Harvest** and **RCP8.5Harvest** compared to reference scenario **BaselineHarvest. B, D**. Latitudinal variation in the spatial *BC*_*ratio*. The shaded areas represented the standard deviation. Latitudes are given in the UTM coordinate system.

Our models predict that beetles would show greater sensitivity to climate variations, given that an increase in dissimilarity was observed even for a medium level of climate change (Figs. 9A), i.e., RCP 4.5 following changes in temperature between the baseline and the two forcing scenarios. In Figs 10A-B, we depicted spatial change in *Tmin* and *WarmMTmean* between RCP8.5 and baseline climate scenarios. *Tmin* and *WarmMTmean* were the most frequently selected respective predictor variables for birds and beetles (see Fig. 6B for the selection percentage). Furthermore, the observed latitudinal dissimilarity gradient was derived mainly from balanced variation in occurrence (*BC*_*ratio* < 0.5 in Figs. 9B, D). Yet, the two taxa behaved in a contrasting manner with regard to their latitudinal variation in the occurrence gradient component of *BC* dissimilarity.

**Figure 10:**
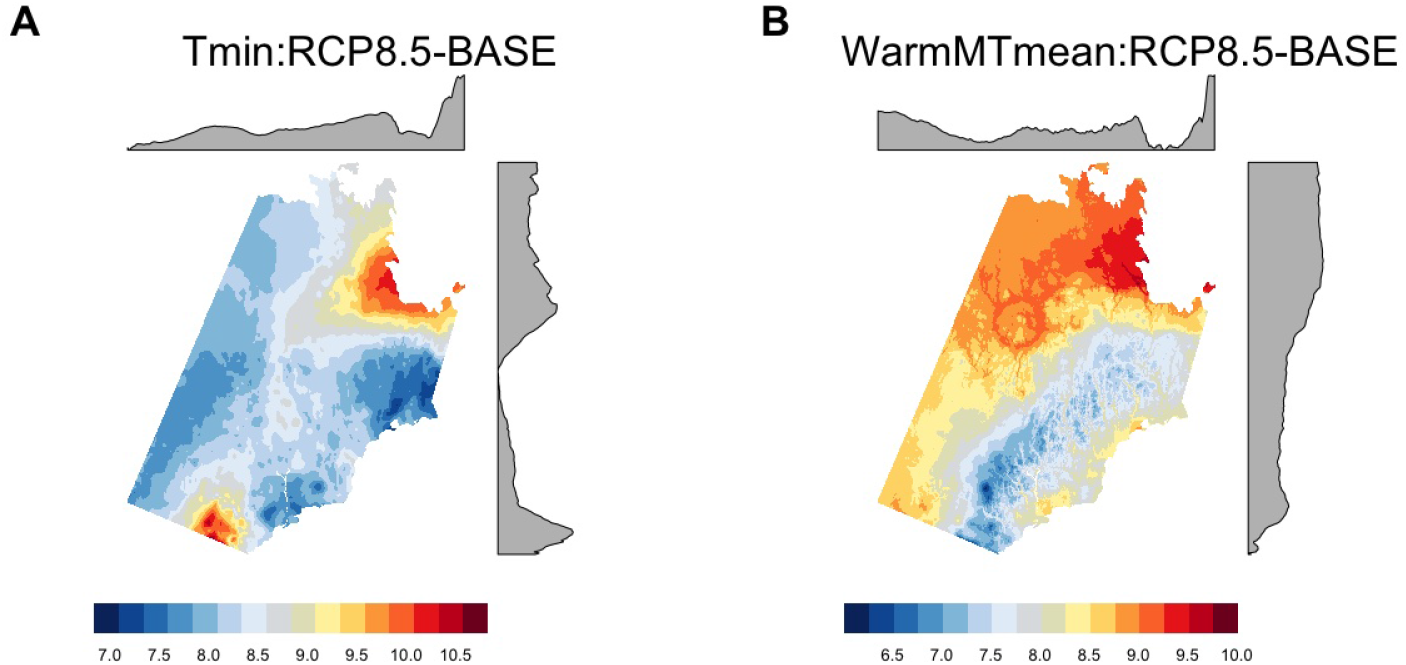
Temperature difference maps between RCP8.5 and baseline of annual minimum temperature (**A**) and mean temperature of the warmest month (**B**) in °C. Map scale: The dark red and the dark blue represented respectively the highest and the lowest values. The horizontal and vertical marginal plots respectively show the longitudinal and latitudinal distribution of the temperature differences.

**Figure 11:**
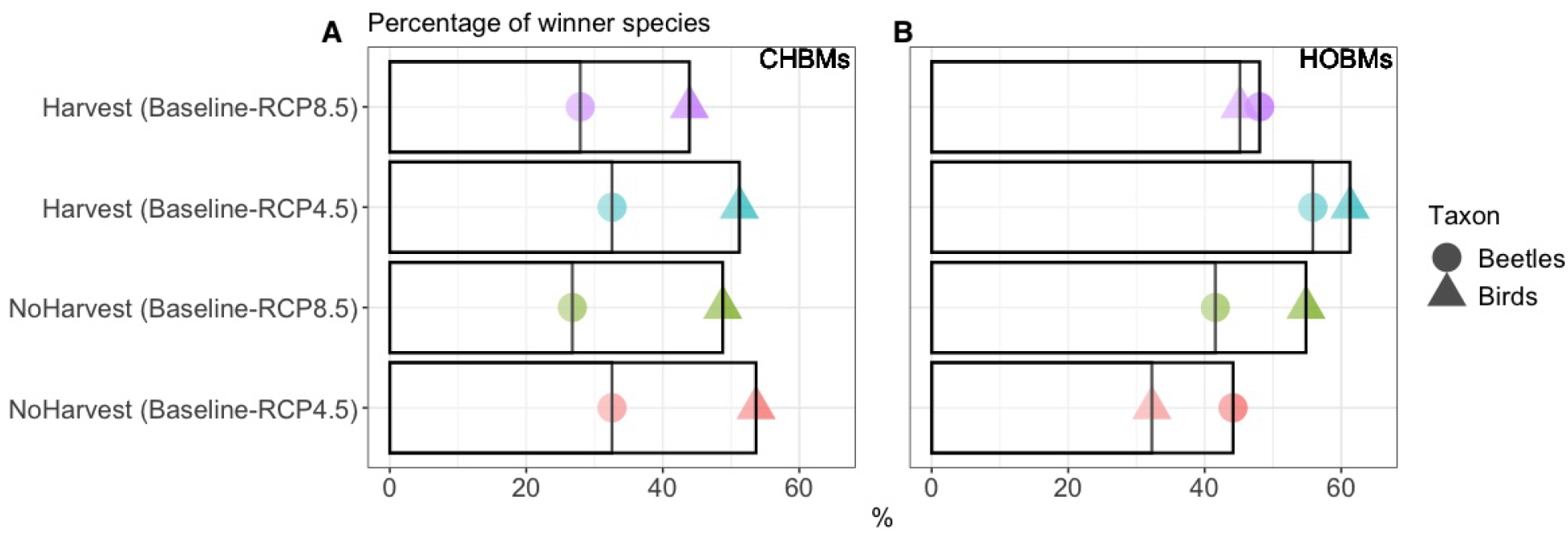
Percentage of winner species per taxon under the four landscapes that were compared and the two model classes (CHBMs and HOBMs). Species *i* was considered a winner if the regional occupancy probability under the simulated scenario was higher than under the reference scenario (*ROP*_*Sim,i*_ > *ROP*_*Ref,i*_).

## 4 Discussion

The present study reveals several intricate effects of global change on animal species assemblages. We showed that the two climate-induced pathways that were studied increased dissimilarity in species assemblage mostly by inducing species turnover, under both forest harvesting scenarios. Of the two pathways, the models integrating both climate and habitat effects strongly modified the assemblage composition more than did the models based only on habitat effects. In fact, the difference in magnitude between the two effects was due to the mismatch that was generated by rapid climate variation compared to slower vegetation change (Wu et al., 2015; Stralberg et al., 2015; Micheletti et al., 2021). Both climate effects caused the decrease in number of winner species from RCP 8.5 to RCP 4.5 compared to the baseline scenario, under the two forest harvesting scenarios. However, some species remained winners under both climate effects such as *Stenichnus turbatus* (Casey) [Staphylindae] and *Chipping Sparrow*, while others changed from winner to loser species, such *Stenichnus perforatus* (Schaum) [Staphylinidae] and *Greater Yellowlegs* (see Table. 6-9 in Supporting Information). Nevertheless, we observed almost an opposite feedback between the two taxa regarding changes in the regional occupancy. This could result from physiological differences between taxa in their individual response to climate.

Our analysis predicts pronounced variations in assemblage composition for both birds and beetles by 2100 that will be caused by climate change under any of the harvest levels. Whereas a rise in anthropogenic radiative forcing should increase dissimilarity in species assemblage following each of the two climate pathways, the magnitude of change differed considerably between the two pathways. For example, the minimum and the maximum assemblage dissimilarities were 0.49 and 0.14 under climate-habitat and habitat-only models respectively. This result could explain the subsequent mismatch between the climate and the biota (Stralberg et al., 2015; Wu et al., 2015; Micheletti et al., 2021). For example, Brice et al. (2020) noted that under climate change the variation in climate conditions would be faster than the capacity of tree species to migrate, a discrepancy that is expected to create a gap between the climatic niche and the observed distribution of species.

Regarding the direction of change at the taxon level, we observed that birds and beetles responded in almost opposite directions. In fact, the increase in bird occupancy that resulted from the mixed combination between direct and indirect climate effects coincided in overall with the *northern biodiversity paradox* that was emphasized by Matthews et al. (2004), Morin and Thuiller (2009), and Berteaux et al. (2010), which anticipated an increase in biodiversity in northern protected areas during this century. More precisely, the northern biodiversity paradox suggests that climate change could lead to a potential increase in the abundance of many species for which the low temperatures are currently a limiting factor (Berteaux et al., 2010). For beetles, we observed a substantial decrease in regional occupancies following an increase in climate change. This outcome aligned with the global scale biodiversity trajectory predicting mostly negative effects of high emissions of greenhouse gas on biodiversity and ecosystem services (Pachauri et al., 2014). Given that beetles are poikilotherms, their internal temperatures vary according to the temperature of the environment, while birds are homeotherms, the internal temperatures of which are physiologically regulated. Our results suggest the potential presence of cold-habitat beetle species in the north. With increasing radiative forcing (Fig. 8F), their occurrence probability decreases as climate conditions extend beyond their tolerance limits. However, those species may also be affected by the competition with southern species invading northern habitats. This could explain the different global implications of direct climate change on beetles and birds. Our conclusions accord with those of other studies based on limitations to phenotypic plasticity and evolvability of critical thermal tolerances in insects (Gaston and Chown, 1999; Terblanche et al., 2007; García-Robledo et al., 2016). For instance, García-Robledo et al. (2016) demonstrated the role of critical thermal temperature on insect tolerance to global warming in the context of elevation, which was used as a proxy for latitude in context of global warming. The authors demonstrated that insects found at middle elevations and on mountain tops were less tolerant to temperature increases than were species located at lower latitudes.

Under mixed climate effects, we observed substantial latitudinal changes in biodiversity with increasing radiative forcing, with an almost complete change in beetle assemblages in the northern portion of the study area. Latitudinal patterns were also noted by Brice et al. (2019), who reported a northward decrease in temporal *Beta* − *diversity* of tree species between past (1970-1980) and present (2000-2016) periods. Our results warn about the possibility that direct climatic effects could increase over the next several decades, particularly for insects. This latitudinal gradient could be a result of the increase in temperature after a change in global climate forcing (Holland and Bitz, 2003). However, the climatic envelope that was used for calibration could increase the effects of the climate change on species occurrence compared to a study that is based upon a greater latitudinal range.

In conclusion, based on implicit assumptions regarding individual species responses to climate change, our analyses identified potential repercussions of two climate-induced pathways that were based on direct and indirect effects on the assemblage composition of two taxonomic groups regionally and latitudinally, under two different forest harvesting scenarios. The observed change was derived mainly by species turnover regardless of harvest level. Moreover, based on the studied species pool and despite the possibility that other species could arrive, we could expect large differences in occupancy between the two studied taxa, which are explained probably by their ecological trait differences. This could indicate the potential range of change in boreal species concerning novel environmental conditions.

## Acknowledgements

This work was supported by the Sentinel North program of Laval University, funded by the Canada First Research Excellence Fund. A.A., D.F., C.H., and P.D. were also supported by the Natural Sciences and Engineering Research Council of Canada (NSERC). We acknowledge Calcul Québec and Compute Canada for their technical support and computing infrastructures. We thank also the Québec Breeding Bird Atlas for supplying data. We would also like to thank the following partners: Regroupement QuébecOiseaux, Environment and Climate Change Canada and Birds Canada, together with all the volunteer participants who gathered data for the project. We are grateful to Louis-Paul Rivest for his valuable statistical suggestions and advice. Special thanks are due to Alexandre Terrigeol for his valuable efforts regarding data collection, Georges Pelletier and Nicolas Bédard for beetle identification and W.F.J. Parsons for his English revision.

