## Supporting Information for "Effects of global change on animal biodiversity in boreal forest landscape: an assemblage dissimilarity analysis"

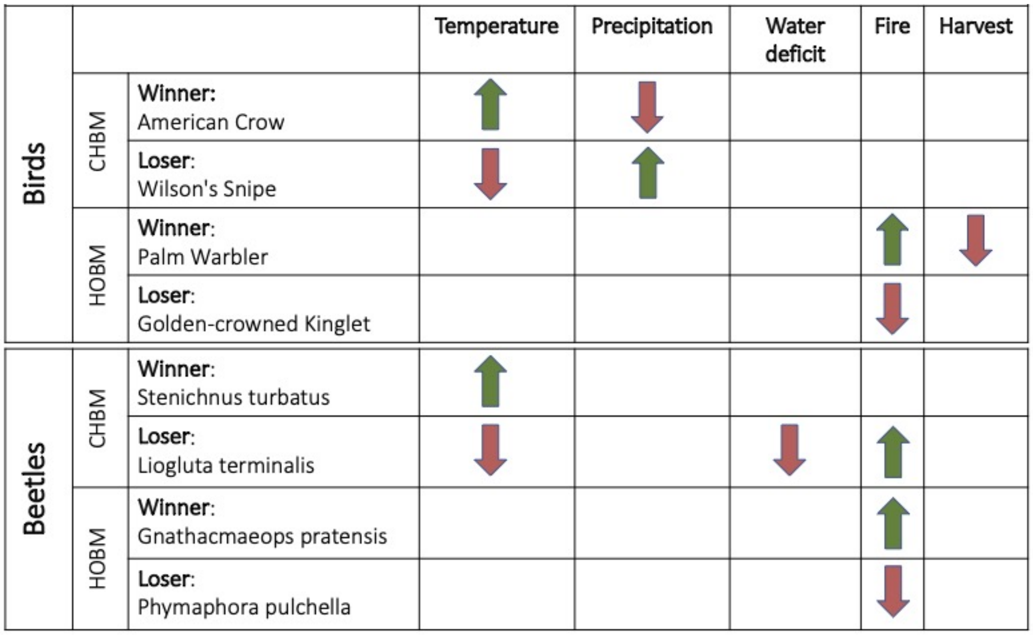


Figure 1: Example of general response of 5 predictor variable classes on 8 winner and loser species.

*

Table 1: Description of the used predictors.

| *Code* | *Variables* |
|---|---|
| Tmax | Annual Maximum Temperature |
| Tmean | Annual Mean Temperature |
| Tmin | Annual Minimum Temperature |
| ColdMTmean | Mean Temperature of Coldest Month |
| ColdQTmean | Mean Temperature of Coldest Quarter |
| WarmMTmean | Mean Temperature of Warmest Month |
| WarmQTmean | Mean Temperature of Warmest Quarter |
| WetQTmean | Mean Temperature of Wettest Quarter |
| DryQTmean | Mean Temperature of Driest Quarter |
| Prcp | Annual Precipitation |
| ColdQPrcp | Precipitation of Coldest Quarter |
| WarmQPrep | Precipitation of Warmest Quarter |
| WetMPrcp | Precipitation of Wettest Month |
| WetQPrcp | Precipitation of Wettest Quarter |
| DryMPrcp | Precipitation of Driest Month |
| DryQPrcp | Precipitation of Driest Quarter |
| WaterDeficit | Evaporation Exceeding the Available Quantity of Available Water |
| ColdQWaterDeficit | WaterDeficit of Coldest Quarter |
| WarmQWaterDeficit | WaterDeficit of Warmest Quarter |
| WetQWaterDeficit | WaterDeficit of Wettest Quarter |
| DryQWaterDeficit | WaterDeficit of Driest Quarter |
| SummerDD5 | Degrees-Day, or Cumulative Temperature Above 0 Degrees. |
| dist_pert_Fire010 | Distance to the Nearest Fire with age in $[0, 10]$ years |
| dist_pert_Fire1020 | Distance to the Nearest Fire with age in $]10, 20]$ years |
| dist_pert_Fire2050 | Distance to the Nearest Fire with age in $]20, 50]$ years |
| Age | Stand Age |
| freqCD | Frequency of Dense Conifer Forests |
| freqCO | Frequency of Open Conifer Forests |
| freqMW | Frequency of Mixed-Wood Habitat |
| freqOH | Frequency of Open Habitat |
| freqDF0_10 | Frequency of Fire disturbed Forests with age in $[0, 10]$ years |
| freqDF10_20 | Frequency of Fire disturbed Forests with age in $]10, 20]$ years |
| freqDF20_50 | Frequency of Fire disturbed Forests with age in $]20, 50]$ years |
| freqDC0_10 | Frequency of Harvest disturbed Forests with age in $[0, 10]$ years |
| freqDC10_20 | Frequency of Harvest disturbed Forests with age in $]10, 20]$ years |
| freqDC20_50 | Frequency of Harvest disturbed Forests with age in $]20, 50]$ years |

Table 2: Natural habitat used (if less than 50% of the pixel's surface is in disturbance).

| Habitat | Condition |
|---|---|
| Dense conifer | - vegetation exceeds more than 50% of the surface<br>- canopy density is higher than 60% of the surface<br>- coniferous stands occupy more than 75% of the surface |
| Open conifer | - vegetation exceeds more than 50% of the surface<br>- canopy density is less than 60% of the surface<br>- coniferous stands occupy more than 75% of the surface |
| Mixed-wood | - vegetation exceeds more than 50% of the surface<br>- neither coniferous nor broad-leaf trees account for more than 75% of the surface<br>- the vegetation surface with trees is greater than the surface without trees |
| Open habitat | - vegetation exceeds more than 50% of the surface<br>- the vegetation surface with trees is less than the surface without trees |

Table 3: The potential 6 full models.

| Models (full) | Fixed-effect without intercept | CHBMs | HOBMs |
|---|---|---|---|
| Model0 | $linear\_terms$ | ✓ | ✓ |
| Model1 | $linear\_terms + Best\_temp : Dist\_Fire\_in + Best\_temp : Age\_in$ | ✓ | × |
| Model2 | $linear\_terms + Age\_in^2$ | ✓ | ✓ |
| Model3 | $linear\_terms + Climate\_in^2$ | ✓ | × |
| Model4 | $linear\_terms + Dist\_Fire\_in^2 + Age\_in^2$ | ✓ | ✓ |
| Model5 | $linear\_terms + Age\_in^2 + Climates\_in^2$ | ✓ | × |

### Table 4: The studied beetle species

| Code | CHBMs | HOBMs | Family | Genus | Species |
|---|---|---|---|---|---|
| CANTPODALAEL | Accepted | Rejected | Cantharidae | Dichelotarsus | laevicollis (Kirby) |
| CARAPTERPUNI | Rejected | Rejected | Carabidae | Pterostichus | punctatissimus (Randall) |
| CARASTERHAEP | Rejected | Rejected | Carabidae | Stereocerus | haematopus Dejean |
| CERATETMCINM | Rejected | Rejected | Cerambycidae | Tetropium | cinnamopterum Kirby |
| CHRYSYNEPILS | Rejected | Rejected | Chrysomelidae | Syneta | pilosa W.J. Brown |
| CLERZENOSANG | Accepted | Accepted | Thanerocleridae | Zenodosus | sanguineus (Say) |
| CRYPCRYPSETS | Accepted | Rejected | Cryptophagidae | Cryptophagus | setulosus Sturm |
| CURCHYLOWARR | Rejected | Rejected | Curculionidae | Hylobius | warreni Wood |
| ELATAMPE | Rejected | Rejected | Elateridae | Ampedus | NA |
| ELATCTENSPIA | Rejected | Rejected | Elateridae | Liotrichus | spinosus (LeConte) |
| ELATCTENTRZO | Rejected | Accepted | Elateridae | Pseudanostirus | triundulatus |
| ELATCTENWATS | Rejected | Accepted | Elateridae | Pseudanostirus | watsoni (W.J. Brown) |
| LATHCORA | Accepted | Accepted | Latridiidae | Corticaria | NA |
| LATHCORRGIBO | Rejected | Rejected | Latridiidae | Cortinicara | gibbosa (Herbst) |
| LATHENICTENO | Accepted | Accepted | Latridiidae | Enicmus | tenuicornis LeConte |
| LYMEELATLUGU | Accepted | Accepted | Lymexylidae | Hylecoetus | lugubris (Say) |
| MELASERRSUBS | Rejected | Rejected | Melandryidae | Serropalpus | substriatus Haldeman |
| MORDMORDBORE | Accepted | Accepted | Mordellidae | Mordellaria | borealis (LeConte) |
| NITIEPURPLAZ | Accepted | Accepted | Nitidulidae | Epuraea | planulata Erichson |
| NITIEPURTERM | Rejected | Rejected | Nitidulidae | Epuraea | terminalis Mannerheim |
| NITIEPURTRUT | Accepted | Accepted | Nitidulidae | Epuraea | truncatella Mannerheim |
| NITIGLISSANS | Accepted | Accepted | Nitidulidae | Glischrochilus | sanguinolentus |
| PTILACRO | Rejected | Rejected | Ptiliidae | Acrotrichis | NA |
| PTILPTIO | Accepted | Accepted | Ptiliidae | Ptiliola | NA |
| RHIZRHIZBRUN | Rejected | Rejected | Monotomidae | Rhizophagus | brunneus |
| RHIZRHIZDIMI | Accepted | Accepted | Monotomidae | Rhizophagus | dimidiatus Mannerheim |
| SCOLDENDRUFP | Rejected | Rejected | Curculionidae | Dendroctonus | rufipennis (Kirby) |
| SCOLDRYOAUTO | Accepted | Accepted | Curculionidae | Dryocoetes | autographus (Ratzeburg) |
| SCOLDRYOBETU | Accepted | Accepted | Curculionidae | Dryocoetes | betulae Hopkins |
| SCOLPOLYRUFP | Accepted | Accepted | Curculionidae | Polygraphus | rufipennis (Kirby) |
| SCOLTRYDLINM | Accepted | Accepted | Curculionidae | Trypodendron | lineatum (Olivier) |
| SCYDPARACOYA | Accepted | Accepted | Staphylinidae | Parascydmus | corpusculus (Casey) |
| STAPATHE | Accepted | Accepted | Staphylinidae | Atheta | NA |
| STAPPHLOLAPP | Accepted | Accepted | Staphylinidae | Phloeostiba | lapponica (Zetterstedt) |
| STAPPHLP | Rejected | Rejected | Staphylinidae | Phloeopora | NA |

| | | | | | |
|---|---|---|---|---|---|
| STAPQUEDLABR | Rejected | Rejected | Staphylinidae | Quedius | labradorensis |
| STAPQUEDPLAG | Accepted | Accepted | Staphylinidae | Quedius | plagiatus Mannerheim |
| STAPSTEH | Accepted | Accepted | Staphylinidae | Stenichnus | NA |
| STAPTACIFRIG | Rejected | Rejected | Staphylinidae | Tachinus | frigidus Erichson |
| TROGTHYMMARQ | Accepted | Accepted | Trogossitidae | Thymalus | marginicollis Chevrolat |
| ANOBHEMICARI | Rejected | Rejected | Ptinidae | Hemicoelus | carinatus (Say) |
| CRYPATOM | Accepted | Accepted | Cryptophagidae | Atomaria | NA |
| CURCHYLOCONG | Rejected | Rejected | Curculionidae | Hylobius | congener Dalla Torre |
| EROTTRIPTHOA | Rejected | Rejected | Erotylidae | Triplax | thoracica Say |
| LATHSTEPBREK | Rejected | Rejected | Latridiidae | Stephostethus | breviclavis (Fall) |
| LYCIDICTAURO | Rejected | Rejected | Lycidae | Dictyoptera | aurora (Herbst) |
| MELAEMMECONN | Rejected | Rejected | Melandryidae | Emmesa | connectens Newman |
| MELAXYLILIVD | Rejected | Rejected | Melandryidae | Dolotarsus | lividus |
| SCOLDRYOAFFR | Accepted | Rejected | Curculionidae | Dryocoetes | affaber (Mannerheim) |
| SCRACANIPALP | Rejected | Rejected | Scraptiidae | Canifa | pallipes (Melsheimer) |
| STAPISCHSPLI | Accepted | Accepted | Staphylinidae | Ischnosoma | splendidum (Gravenhorst) |
| STAPLIOGALAO | Accepted | Accepted | Staphylinidae | Liogluta | aloconotoides Lohse |
| STAPLORDFUNC | Rejected | Rejected | Staphylinidae | Lordithon | fungicola Campbell |
| STAPLYPOFRAM | Accepted | Rejected | Staphylinidae | Lypoglossa | franclemonti Hoebeke |
| STAPOLOPROTL | Accepted | Accepted | Staphylinidae | Olophrum | rotundicolle (C.R. Sahlberg) |
| STAPPLACTACD | Accepted | Rejected | Staphylinidae | Placusa | tachyporoides (Waltl) |
| STAPPLACVAGC | Accepted | Rejected | Staphylinidae | Placusa | vaga Casey |
| STAPQUEDBRUI | Rejected | Rejected | Staphylinidae | Quedius | brunnipennis Mannerheim |
| STAPQUEDRUST | Rejected | Accepted | Staphylinidae | Quedius | rusticus Smetana |
| CARAPTERBREC | Rejected | Rejected | Carabidae | Pterostichus | brevicornis (Kirby) |
| CARATRECCRAC | Accepted | Rejected | Carabidae | Trechus | crassiscapus Lindroth |
| LEIOAGAT | Accepted | Accepted | Leiodidae | Agathidium | NA |
| LEIOLEIO | Accepted | Accepted | Leiodidae | Leiodes | NA |
| PSELEUPL | Accepted | Accepted | Staphylinidae | Euplectus | NA |
| PSELREICSPAL | Rejected | Rejected | Staphylinidae | Reichenbachia | spatulifer Casey |
| STAPACIDQUAR | Accepted | Accepted | Staphylinidae | Acidota | quadrata (Zetterstedt) |
| STAPISCHFIMI | Rejected | Rejected | Staphylinidae | Ischnosoma | fimbriatum Campbell |
| STAPMEGAEXCI | Rejected | Rejected | Staphylinidae | Megarthrus | excisus LeConte |
| STAPPROT | Accepted | Accepted | Staphylinidae | Proteinus | NA |
| STAPQUEDFRIG | Rejected | Rejected | Staphylinidae | Quedius | frigidus Smetana |
| TETRABSTVARG | Accepted | Rejected | Tetratomidae | Tetratoma | variegata Casey |
| CARACALAINGR | Rejected | Rejected | Carabidae | Calathus | ingratus Dejean |
| CRYPCRYPDIFF | Accepted | Accepted | Cryptophagidae | Cryptophagus | difficilis Casey |
| CRYPCRYPSUBF | Accepted | Rejected | Cryptophagidae | Cryptophagus | subfumatus Kraatz |

| | | | | | |
|---|---|---|---|---|---|
| ELATEANUDECO | Rejected | Rejected | Elateridae | Eanus | decoratus (Mannerheim) |
| ELATIDOLDEBI | Rejected | Rejected | Elateridae | Idolus | debilis (LeConte) |
| LATHLATH | Accepted | Rejected | Latridiidae | Latridius | NA |
| LEIOCATO | Rejected | Rejected | Leiodidae | Catops | NA |
| SALPRHINVIRH | Rejected | Rejected | Salpingidae | Rhinosimus | viridiaeneus Randall |
| SCOLPITKSPAR | Accepted | Accepted | Curculionidae | Pityokteines | sparsus (LeConte) |
| SCYDBRACPUBP | Accepted | Accepted | Staphylinidae | Brachycepsis | pubipennis (Casey) |
| STAPLYPOANGR | Accepted | Rejected | Staphylinidae | Lypoglossa | angularis (Mäklin) |
| STAPOXYDGRNP | Rejected | Rejected | Staphylinidae | Oxypoda | sylvia |
| STAPTACIELON | Rejected | Accepted | Staphylinidae | Tachinus | elongatus Gyllenhal |
| CARAPTERADST | Rejected | Accepted | Carabidae | Pterostichus | adstrictus Eschscholtz |
| CIIDORTHPUNC | Accepted | Rejected | Ciidae | Orthocis | punctatus (Mellié) |
| CRYPCRYPCROU | Accepted | Accepted | Cryptophagidae | Cryptophagus | croceus Zimmermann |
| CRYPCRYPPILO | Rejected | Rejected | Cryptophagidae | Cryptophagus | punctipennis |
| LATHCARTCOZV | Rejected | Rejected | Latridiidae | Cartodere | constricta (Gyllenhal) |
| SILPNICUDEFO | Accepted | Rejected | Silphidae | Nicrophorus | defodiens Mannerheim |
| STAPOXYDCOXE | Rejected | Rejected | Staphylinidae | Oxypoda | convergens Casey |
| STAPBRATVARC | Rejected | Rejected | Staphylinidae | Brathinus | varicornis LeConte |
| STAPQUEDFULL | Rejected | Rejected | Staphylinidae | Quedius | fulvicollis (Stephens) |
| CARASPHANITC | Accepted | Rejected | Carabidae | Sphaeroderus | nitidicollis Guérin-Méneville |
| STAPLEPSBREL | Accepted | Accepted | Staphylinidae | Leptusa | brevicollis Casey |
| ELATAGRSLIMO | Rejected | Rejected | Elateridae | Agriotes | limosus (LeConte) |
| LAMPELLYCOZD | Rejected | Rejected | Lampyridae | Ellychnia | corrusca (Linnaeus) |
| LATHMELA | Rejected | Rejected | Latridiidae | Melanophthalma | americana (Mannerheim) |
| PTILPTER | Rejected | Rejected | Ptiliidae | Pteryx | NA |
| SCIRCYPHVARB | Rejected | Accepted | Scirtidae | Cyphon | variabilis (Thunberg) |
| SCOLSCOLPICF | Accepted | Rejected | Curculionidae | Scolytus | piceae (Swaine) |
| CARATRECAPIL | Rejected | Rejected | Carabidae | Trechus | apicalis Motschulsky |
| LEIOAGATEXIS | Accepted | Accepted | Leiodidae | Agathidium | exiguum Melsheimer |
| LEIOANIS | Rejected | Rejected | Leiodidae | Anisotoma | errans W.J. Brown |
| LATHCORNCAVI | Rejected | Rejected | Latridiidae | Corticarina | cavicollis (Mannerheim) |
| PYROISCHCOSA | Rejected | Rejected | Ischaliidae | Ischalia | costata (LeConte) |
| SCYDSTENPEFS | Accepted | Accepted | Staphylinidae | Stenichnus | perforatus (Schaum) |
| STAPPHLOLAED | Rejected | Rejected | Staphylinidae | Phloeonomus | laesicollis (Mäklin) |
| LEIOAGATREPN | Accepted | Accepted | Leiodidae | Agathidium | repentinum Horn |
| ANOBXESTGASN | Rejected | Rejected | Ptinidae | Xestobium | gaspensis R.E. White |
| CUCUDENDCYGN | Rejected | Rejected | Silvanidae | Dendrophagus | cygnaei Mannerheim |
| SILVSILVBIDE | Accepted | Accepted | Silvanidae | Silvanus | bidentatus (Fabricius) |
| EUCNEPIPCORZ | Rejected | Rejected | Eucnemidae | Epiphanis | cornutus Eschscholtz |

| | | | | | |
|---|---|---|---|---|---|
| SCOLSCIEANNT | Rejected | Rejected | Curculionidae | Scierus | annectans LeConte |
| CERAPYGONIGQ | Rejected | Rejected | Cerambycidae | Pygoleptura | nigrella |
| COLYLASCBORE | Rejected | Rejected | Zopheridae | Lasconotus | borealis Horn |
| ELATSERIINCQ | Accepted | Accepted | Elateridae | Sericus | incongruus (LeConte) |
| CLERTHANUNDS | Accepted | Accepted | Cleridae | Thanasimus | undatulus |
| RHIZRHIZREMO | Rejected | Rejected | Monotomidae | Rhizophagus | remotus LeConte |
| STAPPLACINCT | Rejected | Rejected | Staphylinidae | Placusa | incompleta Sjöberg |
| STAPPLACPSUE | Accepted | Accepted | Staphylinidae | Placusa | pseudosuecica Klimaszewski |
| CURCRHYOMACS | Accepted | Accepted | Curculionidae | Rhyncolus | macrops Buchanan |
| CERYCERYCASU | Rejected | Rejected | Cerylonidae | Cerylon | castaneum Say |
| ELATDENTDENC | Rejected | Rejected | Elateridae | Denticollis | denticornis (Kirby) |
| CIIDCISZSTRU | Accepted | Accepted | Ciidae | Cis | striolatus Casey |
| CUCUPEDIFUSC | Accepted | Accepted | Cucujidae | Pediacus | fuscus Erichson |
| NITIEPURLINA | Accepted | Accepted | Nitidulidae | Epuraea | linearis Mäklin |
| SCYDSTENTURT | Accepted | Accepted | Staphylinidae | Stenichnus | turbatus (Casey) |
| STAPGYRP | Accepted | Accepted | Staphylinidae | Gyrophaena | NA |
| STAPSYNTGRAH | Rejected | Accepted | Staphylinidae | Syntomium | grahami Hatch |
| NITIEPURPARN | Accepted | Accepted | Nitidulidae | Epuraea | parsonsi Connell |
| STAPLATH | Rejected | Rejected | Staphylinidae | Lathrobium | elongatum |
| CORYCLYPFUSG | Accepted | Accepted | Corylophidae | Clypastraea | fusca (Harold) |
| ELATEANUESTR | Rejected | Rejected | Elateridae | Eanus | estriatus (LeConte) |
| MELAXYLILAEA | Accepted | Accepted | Melandryidae | Xylita | laevigata (Hellenius) |
| STAPEUCNBRUJ | Accepted | Rejected | Staphylinidae | Eucnecosum | brunnescens (J. Sahlberg) |
| LEIOAGATFAWC | Accepted | Accepted | Leiodidae | Agathidium | fawcettae Miller & Wheeler |
| STAPACIDCREA | Rejected | Rejected | Staphylinidae | Acidota | crenata (Fabricius) |
| STAPACRO | Accepted | Accepted | Staphylinidae | Acrotona | NA |
| CERAACMSPROT | Accepted | Accepted | Cerambycidae | Acmaeops | proteus |
| SALPSPHAVIRE | Rejected | Rejected | Salpingidae | Sphaeriestes | virescens (LeConte) |
| STAPSTEN | Rejected | Rejected | Staphylinidae | Stenus | NA |
| ANOBMICREMAM | Accepted | Rejected | Ptinidae | Microbregma | emarginatum (Duftschmid) |
| CERAGNATPRAT | Accepted | Accepted | Cerambycidae | Gnathacmaeops | pratensis (Laicharting) |
| NITIEPURBORD | Accepted | Accepted | Nitidulidae | Epuraea | rufomarginata |
| SCOLCRYTBORE | Rejected | Rejected | Curculionidae | Crypturgus | borealis Swaine |
| CERAXESTTIBI | Rejected | Rejected | Cerambycidae | Xestoleptura | tibialis (LeConte) |
| CARAPLANDECC | Accepted | Accepted | Carabidae | Platynus | decentis (Say) |
| SCOLPITP | Rejected | Accepted | Curculionidae | Pityophthorus | NA |
| STAPOMALRIVE | Accepted | Accepted | Staphylinidae | Omalium | rivulare (Paykull) |
| CERARHAGINQT | Rejected | Rejected | Cerambycidae | Rhagium | inquisitor (Linnaeus) |

| | | | | | |
|---|---|---|---|---|---|
| NITIEPURAVAR | Rejected | Rejected | Nitidulidae | Epuraea | avara (Randall) |
| CUCULAEMBIGT | Rejected | Rejected | Laemophloeidae | Laemophloeus | biguttatus (Say) |
| PSELPSELBELX | Accepted | Accepted | Staphylinidae | Pselaphus | bellax Casey |
| STAPNUDOCEPU | Rejected | Rejected | Staphylinidae | Nudobius | cephalus (Say) |
| ELATNEOHTUME | Accepted | Accepted | Elateridae | Neohypdonus | tumescens (LeConte) |
| STAPGABIMICQ | Accepted | Accepted | Staphylinidae | Gabrius | microphthalmus (Horn) |
| ENDOPHYMPULE | Accepted | Accepted | Endomychidae | Phymaphora | pulchella Newman |
| STAPDINABORE | Accepted | Accepted | Staphylinidae | Dinaraea | borealis Lohse |
| CARASCAPBILO | Accepted | Rejected | Carabidae | Scaphinotus | bilobus (Say) |
| ELATCTENNITL | Rejected | Rejected | Elateridae | Setasomus | nitidulus (LeConte) |
| MELASERRCOXA | Rejected | Rejected | Melandryidae | Serropalpus | coxalis Mank |
| NITIGLISVITT | Rejected | Accepted | Nitidulidae | Glischrochilus | vittatus (Say) |
| LATHCORALAPP | Accepted | Accepted | Latridiidae | Corticaria | lapponica |
| LATHCORASERR | Accepted | Accepted | Latridiidae | Corticaria | serricollis |
| LATHCORASERT | Accepted | Accepted | Latridiidae | Corticaria | serrata |
| STAPBORSLAMF | Accepted | Accepted | Staphylinidae | Boreostiba | frigida |
| STAPLIOGTERM | Accepted | Accepted | Staphylinidae | Liogluta | terminalis |
| STAPPROTPARQ | Accepted | Accepted | Staphylinidae | Proteinus | parvulus |

### Table 5: The studied bird species

| **Code** | **CHBMs** | **HOBMs** | **Common name** | **Scientific name** |
|---|---|---|---|---|
| ALFL | Accepted | Accepted | Alder Flycatcher | *Empidonax alnorum* |
| AMCR | Accepted | Accepted | American Crow | *Corvus brachyrhynchos* |
| AMGO | Accepted | Accepted | American Goldfinch | *Spinus tristis* |
| AMRE | Accepted | Accepted | American Redstart | *Setophaga ruticilla* |
| AMRO | Rejected | Rejected | American Robin | *Turdus migratorius* |
| BBWA | Accepted | Accepted | Bay-breasted Warbler | *Setophaga castanea* |
| BBWO | Rejected | Rejected | Black-backed Woodpecker | *Picoides arcticus* |
| BCCH | Accepted | Accepted | Black-capped Chickadee | *Poecile atricapillus* |
| BHVI | Accepted | Rejected | Blue-headed Vireo | *Vireo solitarius* |
| BLWA | Accepted | Accepted | Blackpoll Warbler | *Setophaga striata* |
| BOCH | Rejected | Rejected | Boreal Chickadee | *Poecile hudsonicus* |
| BRCR | Accepted | Accepted | Brown Creeper | *Certhia americana* |
| BTNW | Accepted | Rejected | Black-throated Green Warbler | *Setophaga virens* |
| CAGO | Rejected | Accepted | Canada goose | *Branta canadensis* |
| CAJA | Accepted | Rejected | Canada Jay | *Perisoreus canadensis* |
| CEDW | Rejected | Rejected | Cedar Waxwing | *Bombycilla cedrorum* |

| | | | | |
|---|---|---|---|---|
| CHSP | Accepted | Accepted | Chipping Sparrow | *Spizella passerina* |
| CMWA | Accepted | Accepted | Cape May Warbler | *Setophaga tigrina* |
| COLO | Rejected | Rejected | Common Loon | *Gavia immer* |
| CORA | Rejected | Rejected | Common Raven | *Corvus corax* |
| COYE | Accepted | Rejected | Common Yellowthroat | *Geothlypis trichas* |
| CSWA | Accepted | Accepted | Chestnut-sided Warbler | *Setophaga pensylvanica* |
| DEJU | Accepted | Accepted | Dark-eyed Junco | *Junco hyemalis* |
| DOWO | Rejected | Rejected | Downy Woodpecker | *Picoides pubescens* |
| EVGR | Accepted | Accepted | Evening Grosbeak | *Coccothraustes vespertinus* |
| FOSP | Accepted | Accepted | Fox Sparrow | *Passerella iliaca* |
| GCKI | Accepted | Accepted | Golden-crowned Kinglet | *Regulus satrapa* |
| GRYE | Accepted | Accepted | Greater Yellowlegs | *Tringa melanoleuca* |
| HEGU | Accepted | Rejected | Herring Gull | *Larus argentatus* |
| HETH | Rejected | Rejected | Hermit Thrush | *Catharus guttatus* |
| LEFL | Rejected | Rejected | Least Flycatcher | *Empidonax minimus* |
| LISP | Accepted | Accepted | Lincoln's Sparrow | *Melospiza lincolnii* |
| MAWA | Rejected | Accepted | Magnolia Warbler | *Setophaga magnolia* |
| MOWA | Accepted | Rejected | Mourning Warbler | *Geothlypis philadelphia* |
| NAWA | Accepted | Rejected | Nashville Warbler | *Leiothlypis ruficapilla* |
| NOFL | Rejected | Rejected | Northern Flicker | *Colaptes auratus* |
| NOWA | Rejected | Rejected | Northern Waterthrush | *Parkesia noveboracensis* |
| OSFL | Accepted | Rejected | Olive-sided Flycatcher | *Contopus cooperi* |
| OVEN | Accepted | Accepted | Ovenbird | *Seiurus aurocapilla* |
| PAWA | Accepted | Accepted | Palm Warbler | *Setophaga palmarum* |
| PHVI | Accepted | Rejected | Philadelphia Vireo | *Vireo philadelphicus* |
| PISI | Accepted | Rejected | Pine Siskin | *Spinus pinus* |
| PUFI | Rejected | Rejected | Purple Finch | *Haemorhous purpureus* |
| RBNU | Rejected | Rejected | Red-breasted Nuthatch | *Sitta canadensis* |
| RCKI | Accepted | Rejected | Ruby-crowned Kinglet | *Regulus calendula* |
| REVI | Accepted | Accepted | Red-eyed Vireo | *Vireo olivaceus* |
| RUBL | Accepted | Accepted | Rusty Blackbird | *Euphagus carolinus* |
| SASP | Accepted | Accepted | Savannah Sparrow | *Passerculus sandwichensis* |
| SWSP | Rejected | Rejected | Swamp Sparrow | *Melospiza georgiana* |
| SWTH | Accepted | Accepted | Swainson's Thrush | *Catharus ustulatus* |
| TEWA | Accepted | Accepted | Tennessee Warbler | *Leiothlypis peregrina* |
| TRSW | Rejected | Rejected | Tree Swallow | *Tachycineta bicolor* |
| WCSP | Accepted | Accepted | White-crowned Sparrow | *Zonotrichia leucophrys* |

| | | | | |
|---|---|---|---|---|
| WISN | Accepted | Rejected | Wilson's Snipe | *Gallinago delicata* |
| WIWA | Accepted | Accepted | Wilson's Warbler | *Cardellina pusilla* |
| WIWR | Rejected | Rejected | Winter Wren | *Troglodytes hiemalis* |
| WTSP | Accepted | Accepted | White-throated Sparrow | *Zonotrichia albicollis* |
| WWCR | Accepted | Accepted | White-winged Crossbill | *Loxia leucoptera* |
| YBFL | Rejected | Rejected | Yellow-bellied Flycatcher | *Empidonax flaviventris* |
| YBSA | Accepted | Rejected | Yellow-bellied Sapsucker | *Sphyrapicus varius* |
| YEWA | Rejected | Accepted | Yellow Warbler | *Setophaga petechia* |
| YRWA | Rejected | Rejected | Yellow-rumped Warbler | *Setophaga coronata* |

**Table 6: Percentage of change in ROP for beetle species under CHBMs**

| Species | BaseNoHar-RCP4.5NoHar | BaseNoHar-RCP8.5NoHar | BaseHar-RCP4.5Har | BaseHar-RCP8.5Har |
|---|---|---|---|---|
| LATHCORALAPP | -30,48 | -35,48 | -30,48 | -35,48 |
| LATHCORASERR | 8,78 | -46,81 | 8,78 | -46,81 |
| LATHCORASERT | 37,87 | 103220,29 | 362,67 | 124304,51 |
| STAPBORSLAMF | -99,58 | -100,00 | -98,03 | -100,00 |
| STAPPROTPARQ | -2,89 | -23,76 | -4,71 | -10,42 |
| CRYPCRYPCROU | 645,21 | 1263,68 | 296,55 | 519,92 |
| STAPLIOGTERM | -100,00 | -100,00 | -100,00 | -100,00 |
| NITIEPURPLAZ | -6,17 | -19,99 | -4,15 | -11,03 |
| SCYDSTENTURT | 2171,98 | 8465,18 | 2171,98 | 8465,18 |
| STAPDINABORE | -98,13 | -100,00 | -98,13 | -100,00 |
| SCYDSTENPEFS | -99,21 | -100,00 | -99,21 | -100,00 |
| STAPSTEH | -5,73 | 42,57 | -1,37 | 39,59 |
| SCOLPITKSPAR | -100,00 | -100,00 | -100,00 | -100,00 |
| LEIOAGATEXIS | 62,82 | 47,87 | 62,82 | 47,87 |
| RHIZRHIZDIMI | -3,85 | -18,83 | -1,88 | -15,08 |
| LATHCORA | -80,52 | -99,89 | -80,52 | -99,89 |
| PTILPTIO | 26,20 | 14,93 | 23,37 | 22,13 |
| SILVSILVBIDE | -97,53 | -99,98 | -97,53 | -99,98 |
| LEIOAGATFAWC | -14,28 | -57,17 | -24,99 | -44,22 |
| LYMEELATLUGU | -99,86 | -100,00 | -99,86 | -100,00 |
| STAPACRO | 221,47 | 381,56 | 191,13 | 317,65 |
| CRYPATOM | -66,03 | -99,79 | -62,97 | -99,72 |
| CLERZENOSANG | 153,42 | 215,07 | 295,11 | 278,32 |
| STAPGABIMICQ | -93,68 | -99,98 | -94,03 | -99,97 |
| STAPPHLOLAPP | -65,57 | -89,87 | -60,56 | -86,54 |
| ELATNEOHTUME | 312,73 | 549,06 | 350,90 | 511,47 |
| NITIEPURBORD | 4,02 | 13,03 | 1,14 | 17,60 |
| STAPPLACPSUE | -99,55 | -100,00 | -99,64 | -100,00 |
| LEIOAGATREPN | -47,01 | -55,38 | -39,47 | -38,26 |
| STAPATHE | -47,47 | -93,19 | -47,47 | -93,19 |
| CARATRECCRAC | -99,79 | -100,00 | -99,79 | -100,00 |
| CIIDCISZSTRU | -97,70 | -99,85 | -97,49 | -99,81 |
| ANOBMICREMAM | 28,76 | -93,66 | 28,76 | -93,66 |
| CURCRHYOMACS | -100,00 | -100,00 | -100,00 | -100,00 |
| MORDMORDBORE | -2,26 | -39,96 | -9,63 | -24,77 |
| STAPACIDQUAR | -99,64 | -100,00 | -99,62 | -100,00 |
| SCOLDRYOAUTO | -92,91 | -99,98 | -92,91 | -99,98 |

| | | | | |
|---|---|---|---|---|
| **LEIOAGAT** | 82,21 | 146,03 | 85,77 | 153,50 |
| **STAPQUEDPLAG** | -37,41 | -65,59 | -45,78 | -68,76 |
| **SCOLDRYOBETU** | -99,99 | -100,00 | -99,99 | -100,00 |
| **STAPOLOPROTL** | -97,93 | -99,99 | -97,93 | -99,99 |
| **ENDOPHYMPULE** | -97,52 | -100,00 | -97,23 | -100,00 |
| **STAPOMALRIVE** | 17,62 | 27,23 | 17,62 | 27,23 |
| **STAPLIOGALAO** | -96,70 | -99,99 | -96,70 | -99,99 |
| **NITIEPURTRUT** | -67,53 | -98,70 | -69,41 | -98,69 |
| **STAPISCHSPLI** | -88,83 | -99,12 | -86,09 | -98,82 |
| **CRYPCRYPSUBF** | -98,94 | -100,00 | -98,94 | -100,00 |
| **CARASCAPBILO** | 219,97 | 285,73 | 219,97 | 285,73 |
| **STAPLEPSBREL** | 286,72 | 318,76 | 286,72 | 318,76 |
| **STAPGYRP** | 16,44 | 41,25 | 11,37 | 18,09 |
| **CARASPHANITC** | 378,59 | -99,78 | 378,59 | -99,78 |
| **CRYPCRYPDIFF** | 9,03 | 27,60 | 4,43 | 9,21 |
| **ELATSERIINCQ** | -29,29 | -49,90 | -50,94 | -62,59 |
| **NITIEPURPARN** | -97,53 | -99,99 | -97,58 | -100,00 |
| **STAPEUCNBRUJ** | -99,84 | -100,00 | -99,84 | -100,00 |
| **CUCUPEDIFUSC** | -99,89 | -100,00 | -99,94 | -100,00 |
| **SILPNICUDEFO** | 286,29 | 430,44 | 286,29 | 430,44 |
| **TETRABSTVARG** | 264,82 | 282,11 | 264,82 | 282,11 |
| **PSELEUPL** | 25,50 | 38,65 | 37,75 | 59,33 |
| **SCYDBRACPUBP** | -96,43 | -99,97 | -96,43 | -99,97 |
| **CLERTHANUNDS** | 0,52 | -12,02 | 3,73 | -11,38 |
| **TROGTHYMMARQ** | -41,41 | -65,60 | -38,08 | -47,90 |
| **NITIGLISSANS** | -77,56 | -99,70 | -82,28 | -99,74 |
| **SCYDPARACOYA** | -72,34 | -98,79 | -66,02 | -98,59 |
| **CIIDORTHPUNC** | 234,02 | 366,88 | 178,00 | 277,95 |
| **STAPLYPOANGR** | -81,23 | -99,95 | -83,95 | -99,97 |
| **NITIEPURLINA** | 472,57 | 635,85 | 348,77 | 442,86 |
| **CARAPLANDECC** | 0,56 | -17,68 | 4,93 | -13,02 |
| **SCOLTRYDLINM** | -65,12 | -97,80 | -73,19 | -98,13 |
| **LEIOLEIO** | 5,71 | 9,61 | 6,22 | 19,33 |
| **CORYCLYPFUSG** | 1,48 | -4,16 | 2,59 | 9,98 |
| **SCOLSCOLPICF** | 25,16 | 57,26 | -4,26 | -11,80 |
| **SCOLDRYOAFFR** | -85,66 | -99,26 | -87,06 | -99,26 |
| **MELAXYLILAEA** | -79,66 | -99,79 | -85,29 | -99,82 |
| **CANTPODALAEL** | -96,54 | -99,97 | -96,38 | -99,97 |
| **CRYPCRYPSETS** | -97,04 | -99,97 | -97,04 | -99,97 |
| **CERAGNATPRAT** | -81,09 | -93,50 | -81,64 | -95,30 |

| | | | | |
|---|---|---|---|---|
| **STAPPLACTACD** | -95,87 | -100,00 | -95,32 | -100,00 |
| **LATHENICTENO** | -94,55 | -99,92 | -94,55 | -99,92 |
| **CERAACMSPROT** | -99,96 | -100,00 | -99,96 | -100,00 |
| **STAPLYPOFRAM** | -4,43 | -7,09 | -4,96 | -6,73 |
| **SCOLPOLYRUFP** | -0,98 | -8,71 | 0,78 | 0,05 |
| **LATHLATH** | -77,27 | -99,71 | -66,53 | -97,22 |
| **STAPPLACVAGC** | -83,08 | -95,94 | -83,08 | -95,94 |
| **PSELPSELBELX** | -42,57 | -64,27 | -23,61 | -47,03 |
| **STAPPROT** | -60,99 | -97,95 | -62,66 | -97,98 |

#### Table 7: Percentage of change in ROP for bird species under CHBMs

| Species | BaseNoHar-RCP4.5NoHar | BaseNoHar-RCP8.5NoHar | BaseHar-RCP4.5Har | BaseHar-RCP8.5Har |
|---|---|---|---|---|
| WCSP | -100,00 | -100,00 | -100,00 | -100,00 |
| AMGO | 4986,81 | 16244,43 | 5946,11 | 15733,94 |
| AMCR | 12179,37 | 34018,52 | 12179,37 | 34018,52 |
| CHSP | 13026,92 | 43584,87 | 13843,46 | 44046,69 |
| GRYE | -96,98 | -99,76 | -96,98 | -99,76 |
| OVEN | 829,38 | 3262,94 | 868,60 | 3310,46 |
| CSWA | 1368,93 | -20,12 | 1368,93 | -20,12 |
| SASP | -10,65 | -1,94 | -0,87 | -8,85 |
| BCCH | 1331,09 | 6798,01 | 1402,21 | 6080,24 |
| REVI | 537,19 | 639,14 | 508,27 | 476,62 |
| WTSP | -4,28 | 0,10 | -2,21 | -2,17 |
| BLWA | -99,14 | -100,00 | -98,99 | -100,00 |
| WISN | -89,22 | -97,28 | -89,22 | -97,28 |
| CMWA | 222,34 | 1344,72 | 222,34 | 1344,72 |
| FOSP | -96,32 | -99,70 | -95,61 | -99,67 |
| EVGR | 36,25 | 253,31 | 36,25 | 253,31 |
| BBWA | 243,72 | 150,50 | 243,72 | 150,50 |
| DEJU | -74,63 | -98,89 | -74,63 | -98,89 |
| WIWA | -39,85 | -57,70 | -44,47 | -65,63 |
| AMRE | 144,04 | 21,09 | 112,33 | -20,22 |
| RUBL | -75,06 | -98,89 | -83,93 | -99,46 |
| HEGU | 4,26 | 17,38 | 2,02 | 7,62 |
| WWCR | -63,45 | -85,00 | -63,45 | -85,00 |
| YBSA | -94,95 | -100,00 | -95,50 | -100,00 |
| PAWA | 12,75 | 37,69 | -1,13 | 18,97 |
| PISI | -77,65 | -100,00 | -77,65 | -100,00 |
| BRCR | -43,84 | -71,31 | -44,92 | -61,34 |
| OSFL | -20,98 | -94,85 | -30,71 | -97,06 |
| BHVI | 277,67 | 752,00 | 715,70 | 1539,22 |
| MOWA | 260,47 | 706,68 | 268,41 | 728,30 |
| NAWA | 343,26 | 458,18 | 347,33 | 461,75 |
| GCKI | -4,97 | -17,93 | -8,44 | -11,56 |
| RCKI | -6,16 | 26,26 | -6,40 | 28,75 |
| BTNW | 233,91 | 1358,08 | 243,22 | 1676,90 |

| | | | | |
|---|---|---|---|---|
| **ALFL** | 17,97 | 96,75 | 12,03 | 53,69 |
| **PHVI** | 72,35 | -55,84 | 40,71 | -71,96 |
| **TEWA** | 21,20 | 61,52 | 19,80 | 62,64 |
| **COYE** | 137,07 | -93,18 | 111,85 | -95,01 |
| **SWTH** | 18,54 | -51,34 | 16,42 | -51,04 |
| **CAJA** | -74,96 | -98,60 | -74,96 | -98,60 |
| **LISP** | -26,70 | -34,95 | -29,38 | -45,21 |

**Table 8: Percentage of change in ROP for beetle species under HOBMs**

| Species | BaseNoHar-RCP4.5NoHar | BaseNoHar-RCP8.5NoHar | BaseHar-RCP4.5Har | BaseHar-RCP8.5Har |
|---|---|---|---|---|
| LATHCORALAPP | -9,35 | -19,58 | -5,11 | -24,96 |
| CRYPCRYPCROU | -13,18 | -10,44 | -9,83 | -20,17 |
| LATHCORASERR | 3,41 | 6,35 | 0,89 | 8,27 |
| STAPBORSLAMF | -6,36 | -26,63 | -17,97 | -20,97 |
| LATHCORASERT | -26,40 | -54,76 | -20,54 | -30,19 |
| STAPPROTPARQ | -2,89 | -23,76 | -4,71 | -10,42 |
| STAPLIOGTERM | -99,99 | -99,99 | 71,05 | -99,99 |
| STAPDINABORE | -7,75 | -20,24 | -11,09 | -13,04 |
| SCYDSTENTURT | 0,91 | 10,05 | 1,73 | 4,16 |
| SCYDSTENPEFS | 4,50 | 15,56 | 1,16 | 21,36 |
| NITIEPURPLAZ | -4,81 | -12,99 | -3,04 | -6,73 |
| SCOLPITKSPAR | 0,62 | 8,07 | 1,37 | 3,39 |
| STAPSTEH | -6,21 | -13,41 | -2,27 | -16,39 |
| LEIOAGATEXIS | 11,47 | 20,98 | 3,00 | 28,20 |
| RHIZRHIZDIMI | -4,71 | -13,41 | -3,79 | -8,09 |
| SILVSILVBIDE | -4,46 | -10,57 | -2,08 | -13,60 |
| LATHCORA | 0,94 | 1,52 | 0,90 | 1,05 |
| PTILPTIO | -2,64 | -6,63 | -0,34 | -6,32 |
| CRYPATOM | -0,19 | -4,17 | 0,81 | 1,53 |
| LEIOAGATFAWC | -14,28 | -57,17 | -24,99 | -44,22 |
| CLERZENOSANG | -38,90 | -56,79 | -21,77 | -26,44 |
| STAPOMALRIVE | 17,15 | 49,28 | 21,68 | 34,27 |
| LYMEELATLUGU | 1,05 | -14,14 | -1,68 | -7,46 |
| ELATNEOHTUME | 47,99 | 122,92 | 30,49 | 71,06 |
| NITIEPURBORD | 10,19 | 27,06 | 7,58 | 23,05 |
| CURCRHYOMACS | -1,42 | -11,98 | -9,08 | -19,36 |
| LEIOAGATREPN | 0,00 | 0,00 | 5,57 | 6,74 |
| STAPPHLOLAPP | 0,00 | 0,00 | 0,13 | 0,01 |
| STAPACIDQUAR | -4,34 | -17,34 | -8,00 | -7,85 |
| STAPSYNTGRAH | 0,00 | 0,00 | 3,38 | 3,10 |
| STAPATHE | -0,06 | -0,89 | -0,17 | -0,42 |
| STAPACRO | 12,06 | 19,03 | 7,19 | 10,14 |
| MORDMORDBORE | -6,13 | -29,01 | -7,05 | -23,04 |
| CIIDCISZSTRU | 18,97 | 32,68 | 13,07 | 18,32 |
| ENDOPHYMPULE | -40,83 | -51,04 | -28,44 | -29,94 |
| NITIEPURPARN | 11,53 | 25,25 | 9,55 | 2,82 |
| SCOLDRYOAUTO | -3,68 | -11,22 | -2,59 | -5,42 |

| | | | | |
|---|---|---|---|---|
| **STAPLIOGALAO** | -3,74 | -11,95 | -3,96 | -6,38 |
| **STAPGYRP** | 10,24 | 29,20 | 6,11 | 8,89 |
| **NITIEPURLINA** | 11,28 | 31,93 | 5,98 | 11,26 |
| **LEIOAGAT** | -9,58 | -17,48 | 11,07 | -27,72 |
| **STAPQUEDPLAG** | -5,73 | -18,61 | -5,43 | -15,22 |
| **SCOLDRYOBETU** | -6,39 | -13,32 | -3,72 | -4,55 |
| **STAPTACIELON** | 8,04 | 22,17 | 10,88 | 16,40 |
| **ELATSERIINCQ** | 4,17 | 7,50 | 2,98 | 20,30 |
| **CUCUPEDIFUSC** | 9,68 | 13,71 | 13,22 | 24,08 |
| **STAPGABIMICQ** | -7,17 | -30,93 | -19,38 | -26,97 |
| **NITIEPURTRUT** | -0,27 | -4,15 | 0,70 | 1,82 |
| **CARAPLANDECC** | 0,00 | 0,00 | 2,90 | 2,49 |
| **STAPOLOPROTL** | -4,51 | -14,57 | -4,24 | -7,37 |
| **SCIRCYPHVARB** | 0,00 | 0,00 | 3,93 | 3,75 |
| **MELAXYLILAEA** | 10,96 | 20,59 | 4,78 | 26,58 |
| **PSELEUPL** | -10,50 | -14,78 | -6,15 | -7,78 |
| **SCOLTRYDLINM** | -3,83 | -15,17 | -4,92 | -9,10 |
| **SCYDPARACOYA** | 9,41 | 25,65 | 11,26 | 15,42 |
| **SCYDBRACPUBP** | -3,20 | -7,81 | -1,43 | -10,09 |
| **CLERTHANUNDS** | 0,52 | -12,02 | 3,73 | -11,38 |
| **STAPLEPSBREL** | 7,31 | 21,68 | 4,02 | 8,70 |
| **CRYPCRYPDIFF** | 7,42 | 25,64 | 5,12 | 7,02 |
| **STAPISCHSPLI** | 0,53 | -10,95 | 1,97 | -1,52 |
| **CERAGNATPRAT** | 69,23 | 170,62 | 38,70 | 67,84 |
| **STAPPLACPSUE** | -1,69 | 1,58 | 0,01 | -6,57 |
| **CERAACMSPROT** | 11,71 | 41,72 | 14,98 | 25,10 |
| **ELATCTENWATS** | -7,05 | -21,97 | -7,77 | -15,32 |
| **CARAPTERADST** | 4,87 | 10,85 | 2,75 | 4,05 |
| **LEIOLEIO** | 5,71 | 9,61 | 6,22 | 19,33 |
| **SCOLPITP** | 4,76 | 16,34 | 2,64 | 18,18 |
| **CORYCLYPFUSG** | 4,27 | 5,54 | 9,64 | 16,60 |
| **NITIGLISVITT** | 32,41 | 67,72 | 21,59 | 33,37 |
| **SCOLPOLYRUFP** | -2,10 | -9,08 | -0,31 | -2,57 |
| **ELATCTENTRZO** | 3,38 | 5,70 | 5,17 | 6,40 |
| **PSELPSELBELX** | 19,68 | 64,12 | 12,99 | 19,97 |
| **STAPQUEDRUST** | -4,40 | -16,97 | -1,63 | -15,14 |
| **NITIGLISSANS** | -4,62 | -13,82 | -3,56 | -8,96 |
| **LATHENICTENO** | -0,78 | -11,78 | -2,23 | -5,60 |
| **TROGTHYMMARQ** | -42,90 | -59,17 | -35,63 | -40,03 |
| **STAPPROT** | 0,51 | 0,05 | 0,12 | -0,29 |

#### Table 9: Percentage of change in ROP for bird species under HOBMs

| Species | BaseNoHar-RCP4.5NoHar | BaseNoHar-RCP8.5NoHar | BaseHar-RCP4.5Har | BaseHar-RCP8.5Har |
|---|---|---|---|---|
| WCSP | -4,24 | 24,38 | 16,41 | 19,91 |
| GRYE | 89,59 | 194,42 | 47,49 | 82,53 |
| OVEN | -13,33 | -28,82 | 3,06 | -19,12 |
| CHSP | -0,29 | 22,01 | 12,11 | 5,33 |
| AMCR | -9,14 | 3,56 | 0,88 | -6,31 |
| AMGO | -10,21 | 0,85 | 1,34 | -6,69 |
| BLWA | 4,89 | 12,34 | 0,87 | 3,54 |
| SASP | -10,65 | -1,94 | -0,87 | -8,85 |
| REVI | -10,93 | -20,03 | -6,21 | -27,16 |
| WTSP | 2,65 | 8,84 | 1,74 | 2,40 |
| BBWA | -3,66 | -14,98 | -2,79 | -18,20 |
| CMWA | -8,08 | -25,55 | -1,06 | -29,67 |
| CSWA | -3,06 | 15,16 | 5,12 | 29,32 |
| WIWA | 13,64 | 38,84 | 4,24 | 13,16 |
| RUBL | 28,85 | 66,36 | 11,28 | 22,88 |
| BCCH | -4,21 | 12,79 | 0,84 | -4,66 |
| BRCR | -8,88 | -30,12 | -17,70 | -20,48 |
| FOSP | -1,91 | 3,68 | 5,32 | 1,71 |
| AMRE | -7,65 | -15,77 | -9,44 | -22,39 |
| SWTH | -8,46 | -21,85 | -8,89 | -15,90 |
| EVGR | -2,77 | -10,18 | 1,14 | -8,61 |
| PAWA | 25,57 | 57,51 | 11,74 | 45,35 |
| WWCR | -4,71 | -23,44 | -4,82 | -12,43 |
| ALFL | -2,02 | -4,70 | -2,67 | -5,88 |
| CAGO | 6,50 | 15,93 | 3,96 | 7,41 |
| GCKI | -8,08 | -25,56 | -11,80 | -17,26 |
| LISP | 10,20 | 28,92 | 4,35 | 6,74 |
| TEWA | -4,52 | -6,32 | -3,76 | -8,86 |
| DEJU | 2,04 | 3,29 | 2,40 | 6,38 |
| MAWA | -6,85 | -17,80 | -8,53 | -14,30 |
| YEWA | 16,51 | 44,68 | 10,20 | 23,59 |
